## Supplementary figures for "Structural Basis of CSN-mediated SCF Deneddylation"

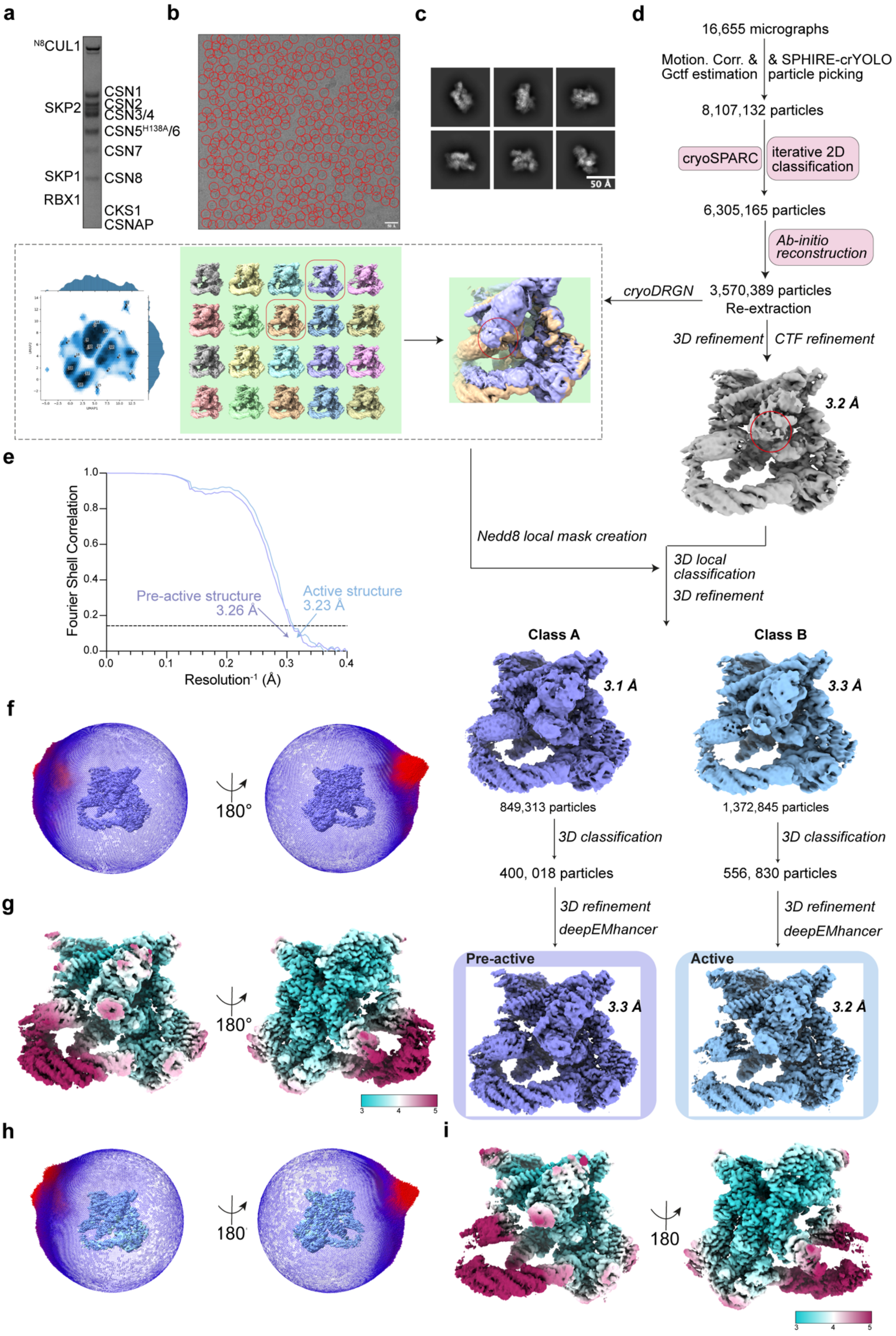
a
N8CUL1
SKP2
SKP1
RBX1
CSN1
CSN2
CSN3/4
CSN5H138A/6
CSN7
CSN8
CKS1
CSNAP
b
c
50 Å
d
16,655 micrographs
Motion. Corr. & Gctf estimation
& SPHIRE-crYOLO particle picking
8,107,132 particles
cryoSPARC
iterative 2D classification
6,305,165 particles
Ab-initio reconstruction
cryoDRGN
3,570,389 particles
Re-extraction
3D refinement
CTF refinement
3.2 Å
Nedd8 local mask creation
3D local classification
3D refinement
Class A
Class B
3.1 Å
3.3 Å
849,313 particles
1,372,845 particles
3D classification
400, 018 particles
556, 830 particles
3D refinement
deepEMhancer
Pre-active
Active
3.3 Å
3.2 Å
e
Fourier Shell Correlation
Resolution-1 (Å)
Pre-active structure 3.26 Å
Active structure 3.23 Å
f
180°
g
180°
h
180
i
180

**Extended Data Fig. 1. Reconstitution, cryo-electron microscopy, and single particle analysis of CSN$^{5H138A}$-$^{N8}$SCF.**

**a,** Coomassie-stained SDS PAGE of the purified CSN$^{5H138A}$-$^{N8}$SCF protein complex. **b,** A representative motion-corrected micrograph and particle picking (red circles). **c,** Representative high-quality 2D reference-free 2D class averages. **d,** Single particle analysis workflow for the CSN-$^{N8}$SCF dataset, resulting in 3D reconstructions of pre- and activated CSN$^{5H138A}$-$^{N8}$SCF. 3D local classification was performed in RELION-4.0 (Kimanius, Dong et al. 2021) utilising a NEDD8-specific mask (generated from classification results from CryoDRGN (Zhong, Bepler et al. 2021) ). **e,** Resolution estimates of pre-activated and activated CSN$^{5H138A}$-$^{N8}$SCF. **f,** Euler angle distribution plots and (**g**) local resolution estimates for pre-activated CSN$^{5H138A}$-$^{N8}$SCF. **h,** Euler angle distribution plots and (**i**) for activated CSN$^{5H138A}$-$^{N8}$SCF. Resolutions for all maps in this figure were estimated using the gold-standard FSC 0.143 criterion.

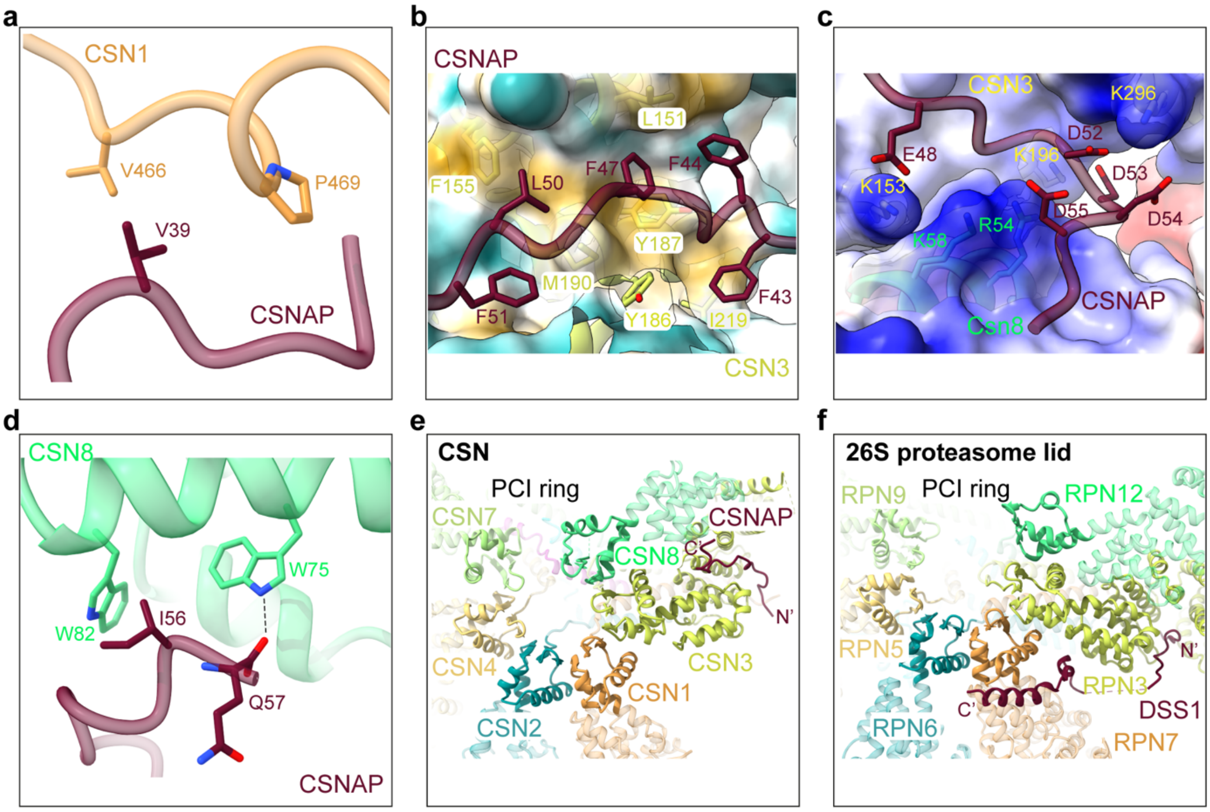


**Extended Data Fig. 2. CSNAP is an integral subunit of the CSN complex.**

**a,** Structural interactions between CSN1 and CSNAP. **b,** CSN3 engages CSNAP through hydrophobic interactions. CSN3 is displayed as a hydrophobic surface, with hydrophobicity mapped using a colour scale (yellow denoting hydrophobic residues and cyan for hydrophilic residues). **c,** An electropositive groove formed by CSN3 and CSN8 accommodates the acidic tail of CSNAP. The surfaces of CSN3 and CSN8 are coloured by electrostatic potential. **d,** CSN8 secures the C-terminus of CSNAP. **e-f,** Structural comparison between CSNAP in CSN (**e**) and DSS1, a subunit of the 26S proteasome lid complex (**f**) (PDB: 3JCK). The subunit colour code for the 26S proteasome lid corresponds to CSN.

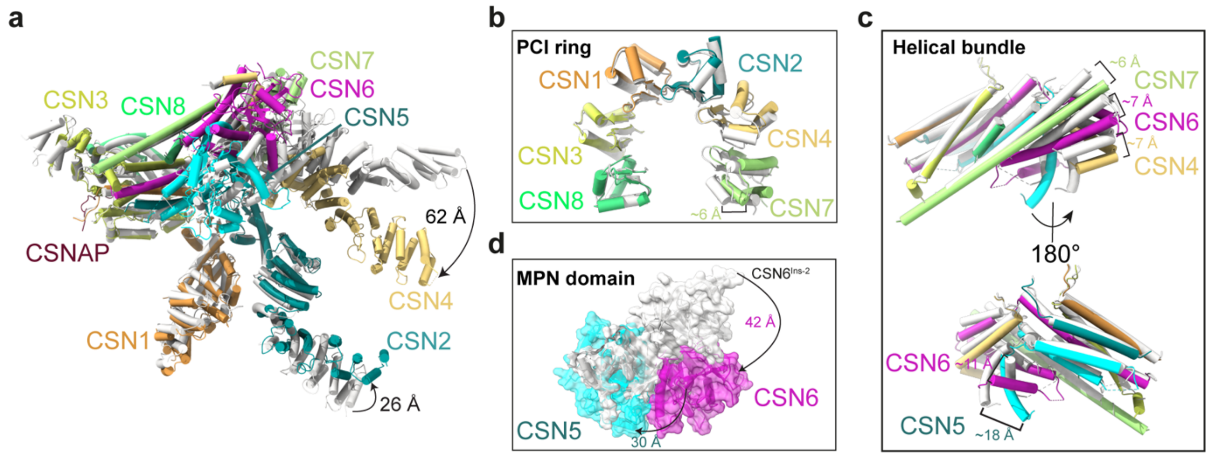


**Extended Data Fig. 3. Conformational changes in CSN between the CSN$^{apo}$ and pre-activated CSN$^{5H138A}$ states.**

**a,** Overall conformational changes of the CSN complex between CSN$^{apo}$ (PDB: 4D10) (grey) and pre-activated CSN$^{5H138A}$-$^{N8}$SCF (coloured). The structural alignment was performed using CSN3 as a reference. **b,** Comparison of the CSN PCI ring architecture between CSN$^{apo}$ (grey) and pre-activated CSN$^{5H138A}$-$^{N8}$SCF (coloured). **c,** Structural comparison of the CSN C-terminal helical bundle. **d,** Close-up view of the MPN domains of CSN5 and CSN6, illustrating conformational changes essential for CSN activation.

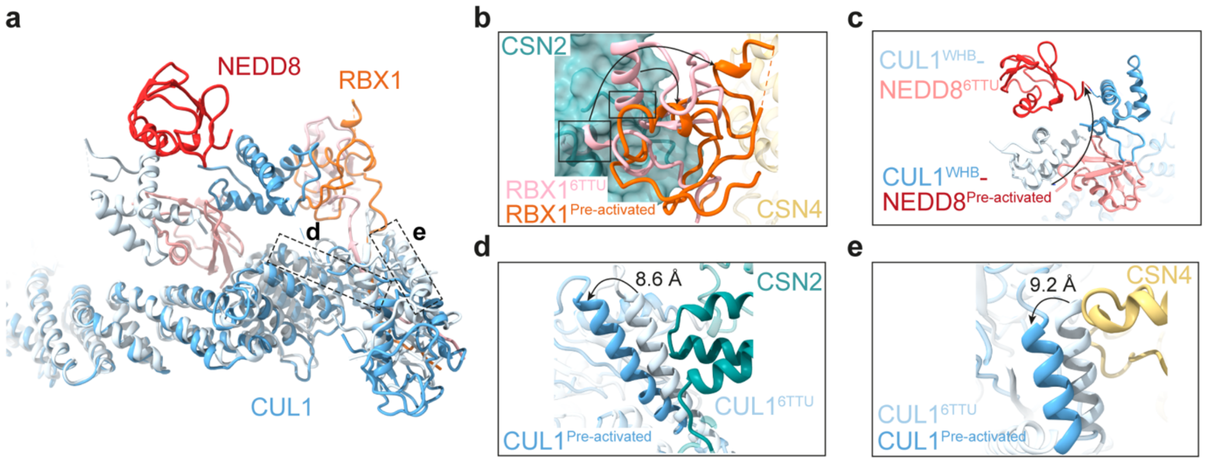


**Extended Data Fig. 4. Conformational changes in $^{N8}$SCF between an active $^{N8}$SCF ubiquitin-bound state and the pre-activated CSN$^{5H138A}$ complex.**

**a,** Overall structural comparison of $^{N8}$SCF between the active UBE2D2-$^{N8}$SCF complex (PDB: 6TTU) (grey) and pre-activated CSN$^{5H138A}$-$^{N8}$SCF (coloured) highlighting key conformational shifts. **b,** Rotation of RBX1$^{RING}$ from its position in UBE2D2-$^{N8}$SCF (grey) to its new position in pre-activated CSN$^{5H138A}$-$^{N8}$SCF (coloured), preventing steric clashes with CSN2. **c,** Close-up view of conformational rearrangements in $^{N8}$WHB. **d,** Structural adjustments in CUL1 to avoid a clash with CSN2. **e,** Additional conformational changes in CUL1 to avoid a clash with CSN4.

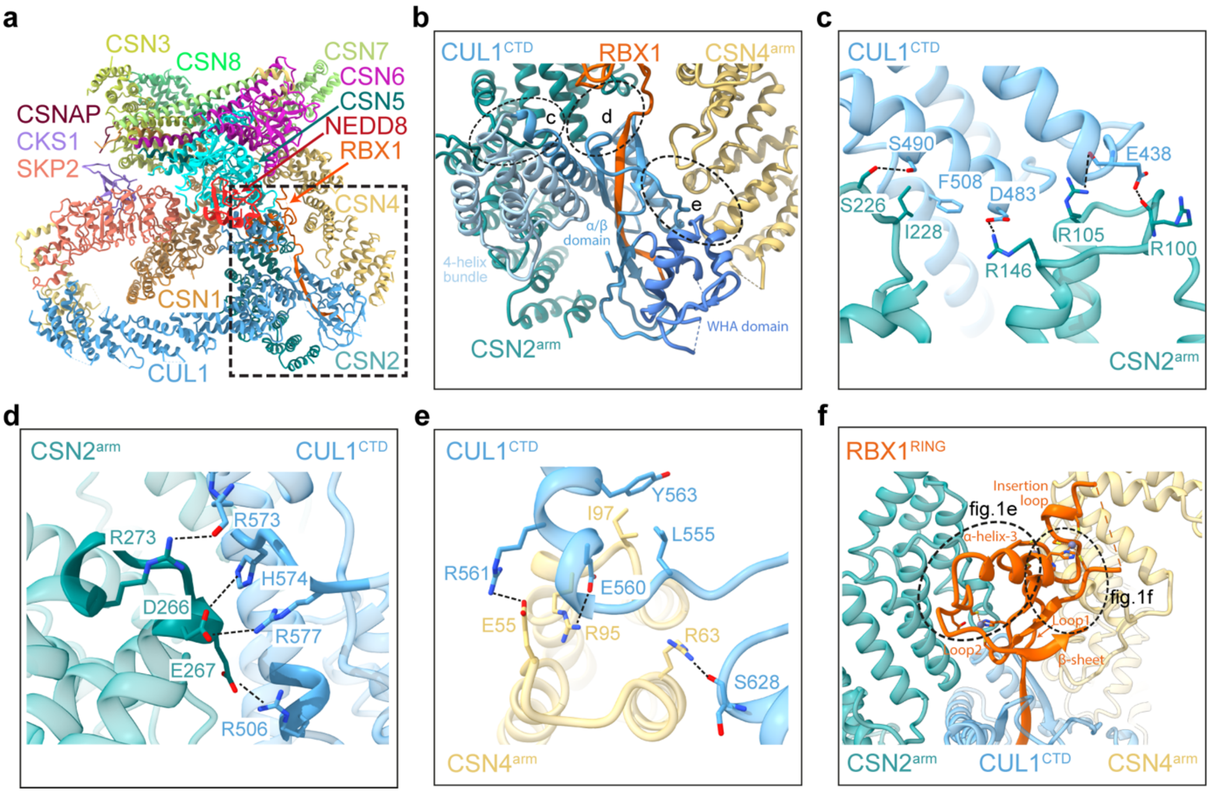


**Extended Data Fig. 5. CSN2$^{arm}$ and CSN4$^{arm}$ clamp CUL1$^{CTD}$ and stabilise RBX1$^{RING}$ in pre-activated CSN$^{5H138A}$-$^{N8}$SCF.**

**a,** Molecular model of pre-activated CSN$^{5H138A}$-$^{N8}$SCF. **b,** Close-up view of the boxed region in (**a**), illustrating how CSN2$^{arm}$ and CSN4$^{arm}$ clamp the CUL1$^{CTD}$ at three distinct regions (highlighted by circles with further details in panels **c**, **d** and **e**). The 4-helix bundle (4-HB), α/β and WHA subdomains of CUL1$^{CTD}$ are coloured in shades of blue. **c,** Detailed view of interface “c”, illustrating CSN2$^{arm}$ interactions with CUL1$^{CTD}$. **d,** Detailed view of interface “d”, illustrating CSN2$^{arm}$ interactions with CUL1$^{CTD}$. **e,** Detailed view of interface “e”, showing CSN4$^{arm}$ interactions with CUL1$^{CTD}$. **f,** Close-up view of the boxed region in (**a**), showing the RBX1$^{RING}$ stabilised by CSN2$^{arm}$ and CSN4$^{arm}$. Dashed circles highlight interaction interfaces related to Fig.1e and Fig.1f.

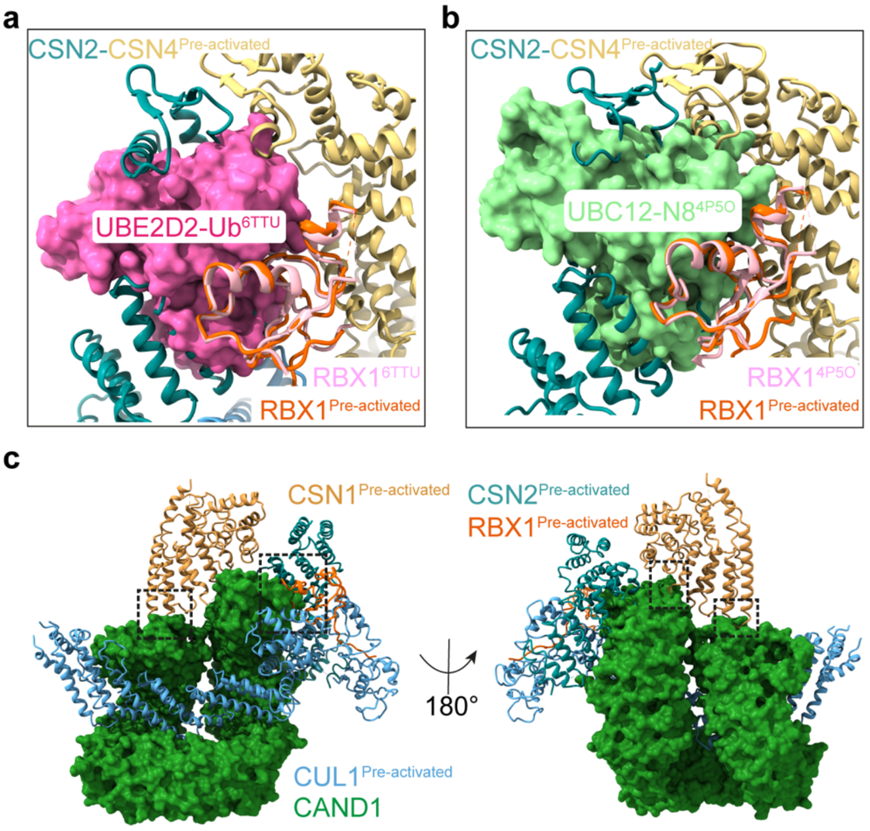


**Extended Data Fig. 6. CSN directly competes with key components of the CRL regulatory cycle.**

**a,** Structural overlay of the ubiquitin E2 (pink surface) from active UBE2D2-$^{N8}$SCF complex (PDB: 6TTU) with pre-activated CSN$^{5H138A}$-$^{N8}$SCF illustrating a steric clash that prevents simultaneous E2 and CSN binding. **b,** Overlay of NEDD8 E2 (green surface) from the RBX1-UBC12~$^{N8}$CUL1-DCNL1 complex (PDB: 4P5O) with pre-activated CSN$^{5H138A}$-$^{N8}$SCF. **c,** Overlay of CAND1 (dark green surface) from the CAND1-CUL1/RBX1-SKP1/SKP2/CKS1-CDK2 complex (PDB: 8OR0) with pre-activated CSN$^{5H138A}$-$^{N8}$SCF, revealing steric clashes between CSN1, CSN2 and RBX1 with CAND1 (highlighted by black boxes).

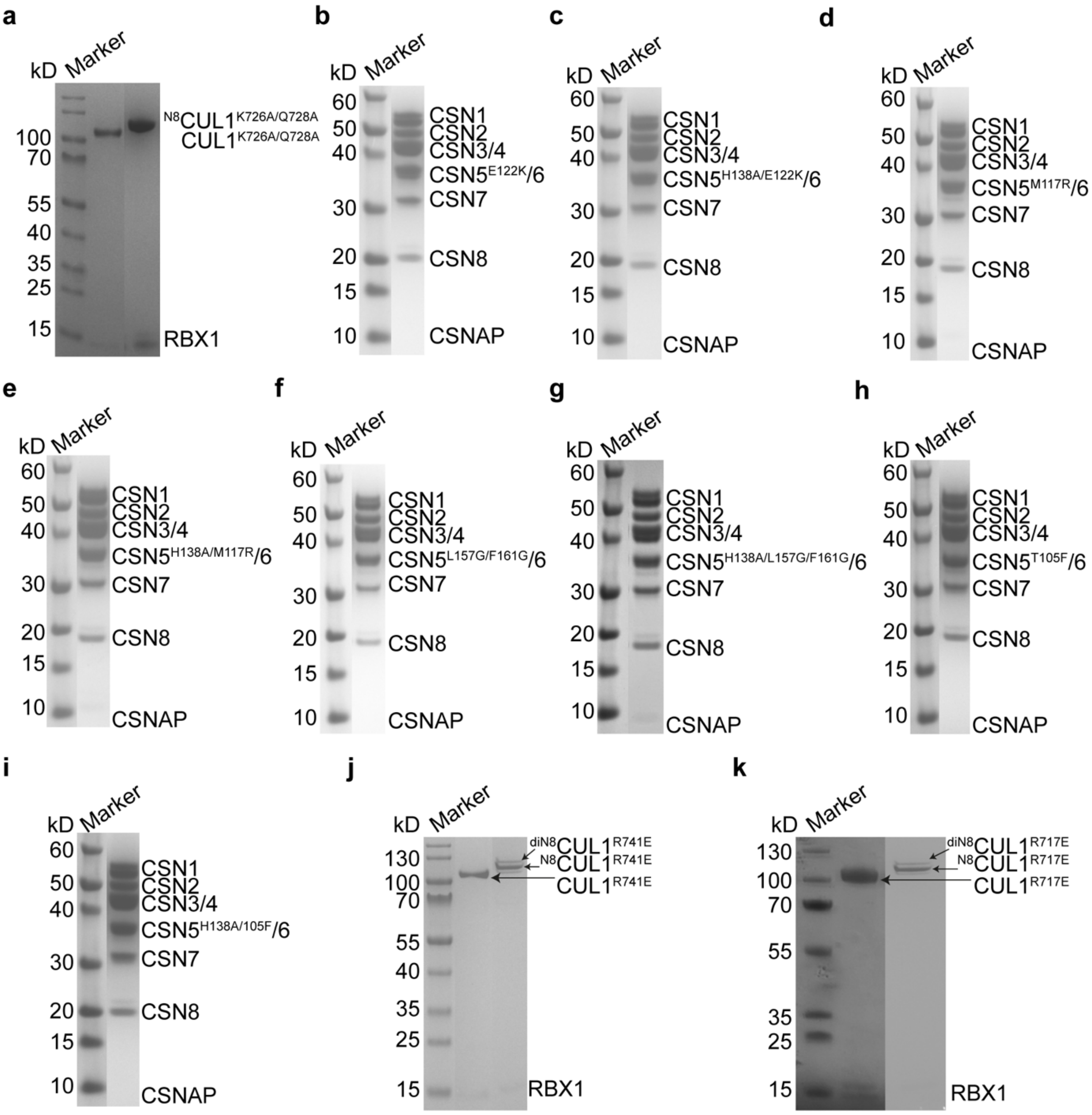


**Extended Data Fig. 7. SDS-PAGE (4-12%) analysis and Coomassie staining of purified mutants.**

**a,** Non-neddylated and neddylated $CUL1^{K726A/Q728A}$/RBX1**. b,** $CSN^{5E122K}$. **c,** $CSN^{5H138A/E122K}$. **d,** $CSN^{5M117R}$. **e,** $CSN^{5H138A/E122K}$. **f,** $CSN^{5L157G/F161G}$. **g,** $CSN^{5H138A/L157G/F161G}$. **h,** $CSN^{5T105F}$. **i,** $CSN^{5H138A/T105F}$. **j,** Non-neddylated and neddylated $CUL1^{R741E}$/RBX1**. k,** Non-neddylated and neddylated $CUL1^{R717E}$/RBX1.

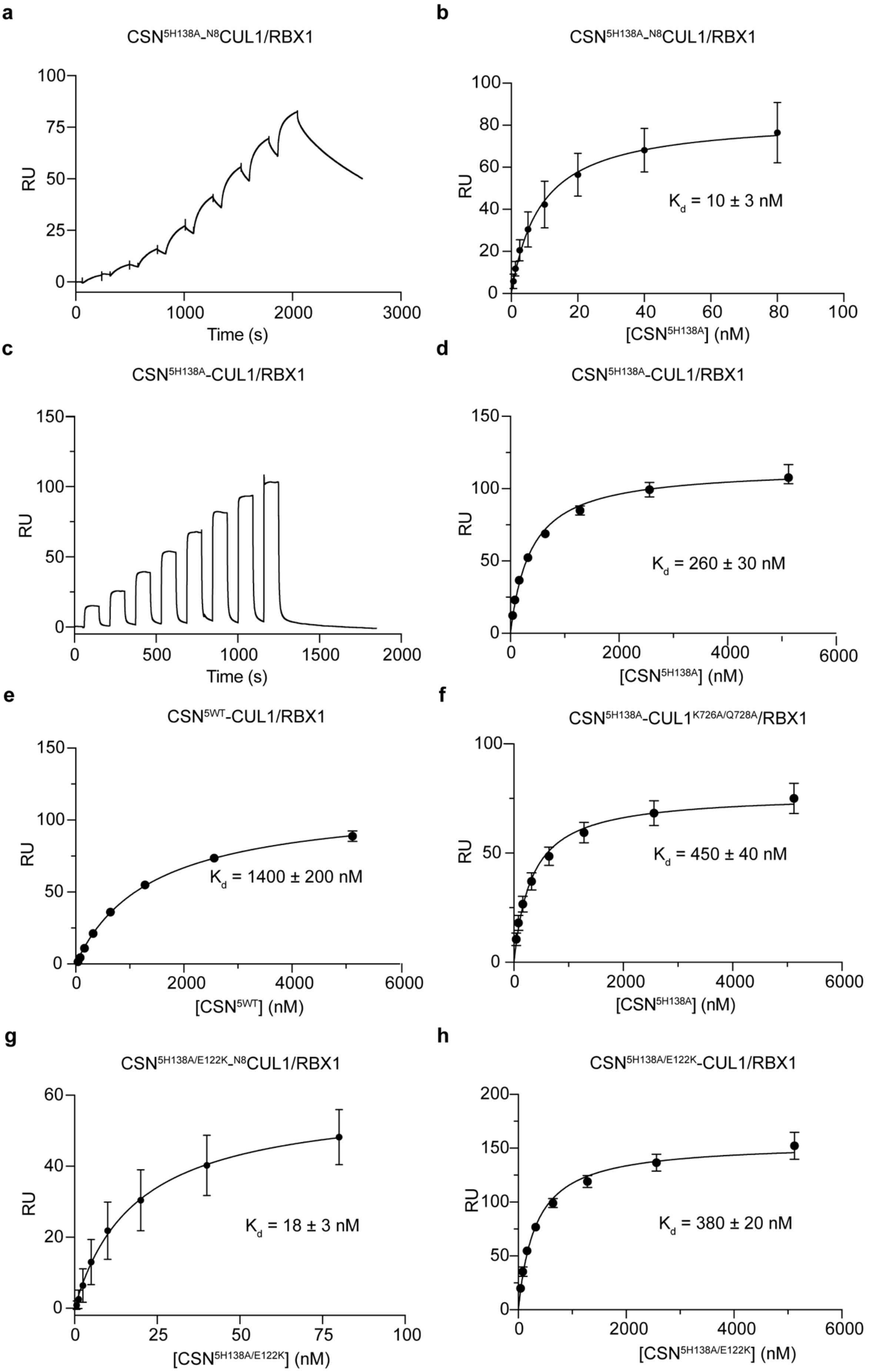
a
CSN5H138A-N8CUL1/RBX1
RU
Time (s)
b
CSN5H138A-N8CUL1/RBX1
RU
Kd = 10 ± 3 nM
[CSN5H138A] (nM)
c
CSN5H138A-CUL1/RBX1
RU
Time (s)
d
CSN5H138A-CUL1/RBX1
RU
Kd = 260 ± 30 nM
[CSN5H138A] (nM)
e
CSN5WT-CUL1/RBX1
RU
Kd = 1400 ± 200 nM
[CSN5WT] (nM)
f
CSN5H138A-CUL1K726A/Q728A/RBX1
RU
Kd = 450 ± 40 nM
[CSN5H138A] (nM)
g
CSN5H138A/E122K-N8CUL1/RBX1
RU
Kd = 18 ± 3 nM
[CSN5H138A/E122K] (nM)
h
CSN5H138A/E122K-CUL1/RBX1
RU
Kd = 380 ± 20 nM
[CSN5H138A/E122K] (nM)

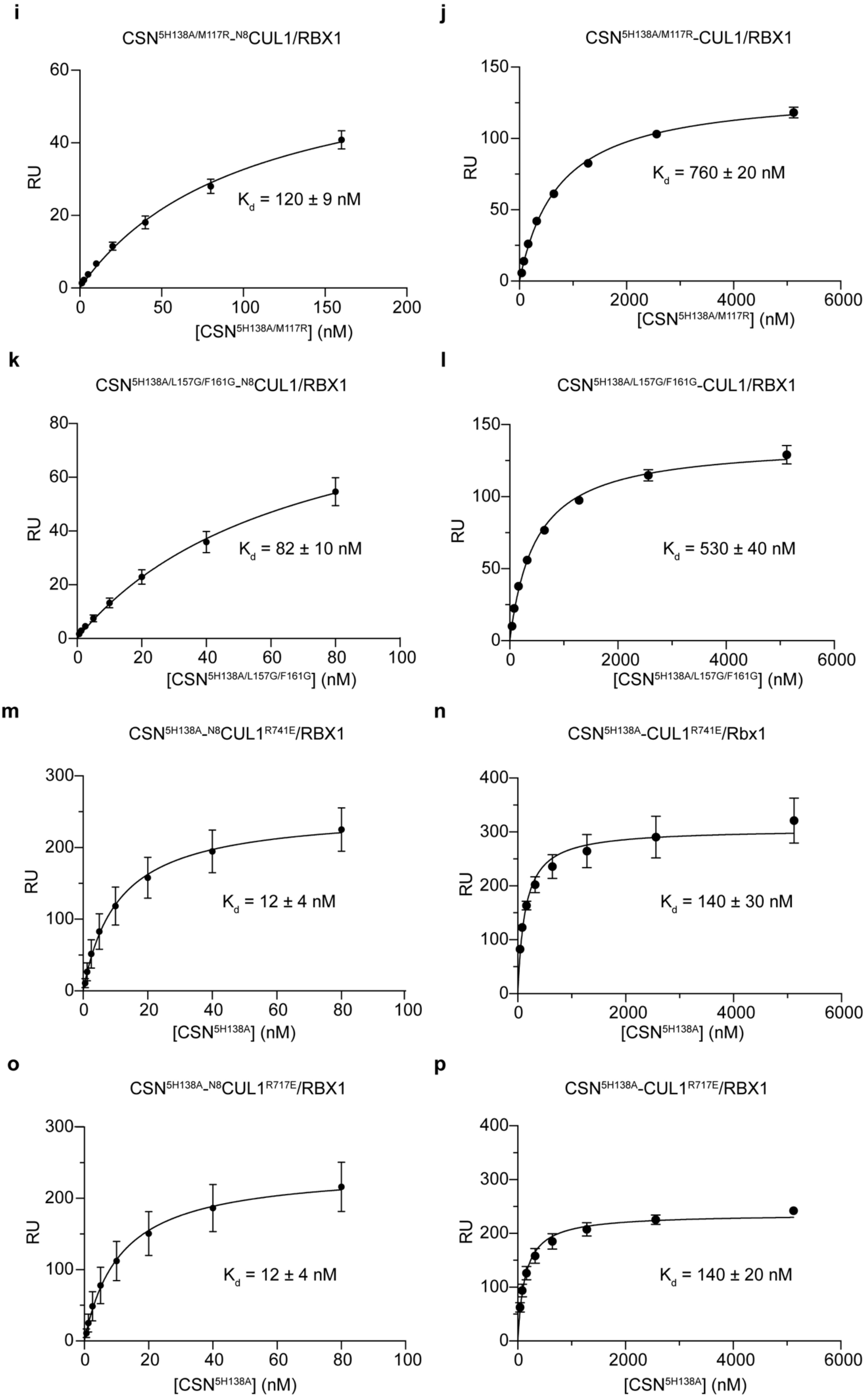
i
CSN5H138A/M117R_N8CUL1/RBX1
RU
Kd = 120 ± 9 nM
[CSN5H138A/M117R] (nM)
j
CSN5H138A/M117R-CUL1/RBX1
RU
Kd = 760 ± 20 nM
[CSN5H138A/M117R] (nM)
k
CSN5H138A/L157G/F161G_N8CUL1/RBX1
RU
Kd = 82 ± 10 nM
[CSN5H138A/L157G/F161G] (nM)
l
CSN5H138A/L157G/F161G-CUL1/RBX1
RU
Kd = 530 ± 40 nM
[CSN5H138A/L157G/F161G] (nM)
m
CSN5H138A_N8CUL1R741E/RBX1
RU
Kd = 12 ± 4 nM
[CSN5H138A] (nM)
n
CSN5H138A-CUL1R741E/Rbx1
RU
Kd = 140 ± 30 nM
[CSN5H138A] (nM)
o
CSN5H138A_N8CUL1R717E/RBX1
RU
Kd = 12 ± 4 nM
[CSN5H138A] (nM)
p
CSN5H138A-CUL1R717E/RBX1
RU
Kd = 140 ± 20 nM
[CSN5H138A] (nM)

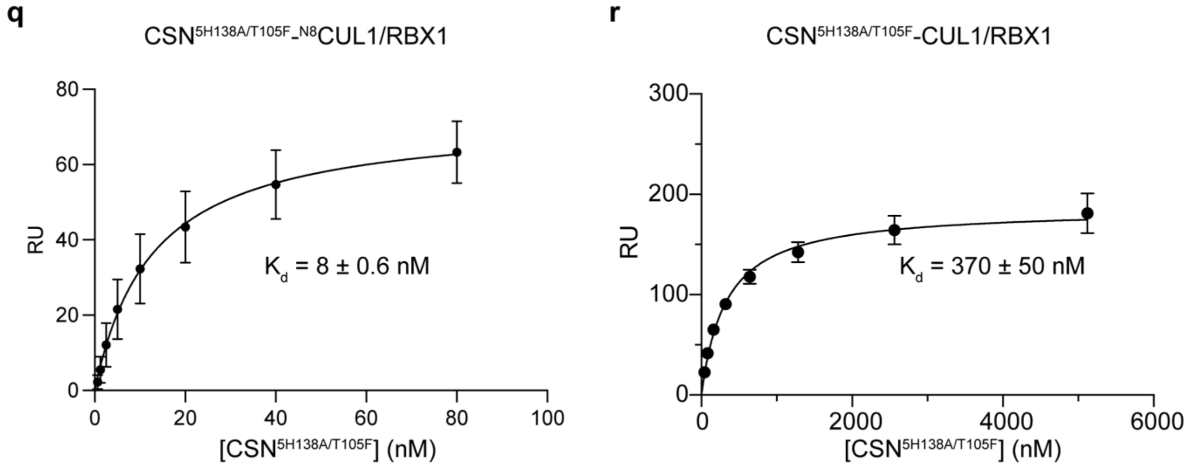


**Extended Data Fig. 8. SPR Analysis of Binding Affinities between CUL1/Rbx1 and CSN mutants.**

**a,** SPR Sensogram of CSN$^{5H138A}$ binding to $^{\text{StrepII2x-N8}}$CUL1/RBX1. **b,** Steady-state binding curve of CSN$^{5H138A}$ to $^{\text{StrepII2x-N8}}$CUL1/RBX1 determined from data in (**a**). The continuous line shows the result of a hyperbolic curve fit to determine $K_d$**. c,** SPR Sensogram of CSN$^{5H138A}$ binding to $^{\text{StrepII2x}}$-CUL1/RBX1. **d,** Steady-state binding curve of CSN$^{5H138A}$ to $^{\text{StrepII2x-N8}}$CUL1/RBX1 determined from data in (**c**). **e,** Binding curve of CSN$^{5WT}$ and $^{\text{StrepII2x}}$-CUL1/RBX1. **f,** Binding curve for affinity determination of CSN$^{5H138A}$ to $^{\text{StrepII2x-N8}}$CUL1$^{K726A/Q728A}$/RBX1. **g,** Binding curve of CSN$^{5H138A/E122K}$ and $^{\text{StrepII2x-N8}}$CUL1/RBX1. **h,** Binding curve of CSN$^{5H138A/E122K}$ and $^{\text{StrepII2x}}$-CUL1/RBX1. **i,** Binding curve of CSN$^{5H138A/M117R}$ and $^{\text{StrepII2x- N8}}$CUL1/RBX1. **j,** Binding curve of CSN$^{5H138A/M117R}$ and $^{\text{StrepII2x}}$- CUL1/RBX1. **k,** Binding curve of CSN$^{5H138A/L157G/F161G}$ and $^{\text{StrepII2x- N8}}$CUL1/RBX1. **l,** Binding curve of CSN$^{5H138A/L157G/F161G}$ and $^{\text{StrepII2x}}$-CUL1/RBX1. **m,** Binding curve of CSN$^{5H138A}$ and $^{\text{StrepII2x- N8}}$CUL1$^{R741E}$/RBX1. **n,** Binding curve of CSN$^{5H138A}$ and $^{\text{StrepII2x}}$- CUL1$^{R741E}$/RBX1. **o,** Binding curve of CSN$^{5H138A}$ and $^{\text{StrepII2x- N8}}$CUL1$^{R717E}$/RBX1. **p,** Binding curve of CSN$^{5H138A}$ and $^{\text{StrepII2x}}$-CUL1$^{R717E}$/RBX1. **q,** Binding curve of CSN$^{5H138A/T105F}$ and $^{\text{StrepII2x- N8}}$CUL1/RBX1. **r,** Binding curve of CSN$^{5H138A/T105F}$ and $^{\text{StrepII2x}}$- CUL1/RBX1. Data represent the mean ± SD from three independent experiments. Dissociation constants were determined using a one-site binding hyperbola.

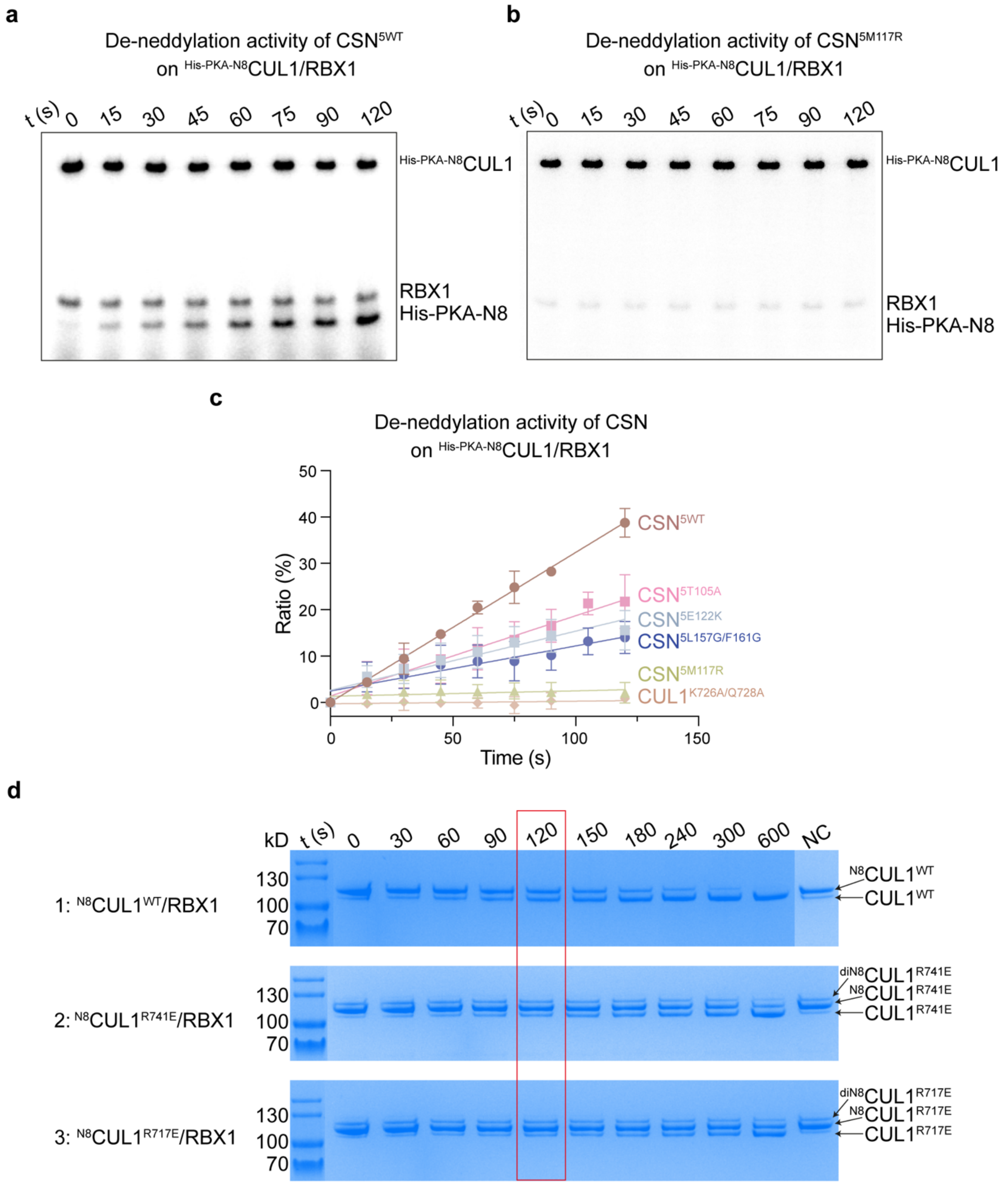


**Extended Data Fig. 9. *In vitro* De-neddylation activity assays.**

**a, b,** SDS-PAGE (4-12%) analysis coupled with phosphate radioactive imaging of *in vitro* de-neddylation assays comparing CSN$^{5WT}$ (**a**) and CSN$^{5M117R}$ (**b**) on $^{His\text{-}PKA\text{-}N8}$CUL1/RBX1. **c,** Quantification of de-neddylation activity measured as the increasing ratio of free NEDD8 to total NEDD8 (free NEDD8 + CUL1-bound NEDD8) over time. The de-neddylation activity is indicated by the slope of the linear regression curve, with mutations in CSN or CUL1 distinguished by colour-coded data points. **d,** SDS-PAGE (4-12%) and Coomassie staining of *in vitro* de-neddylation assays comparing 1: $^{N8}$CUL1/RBX1; 2: $^{N8}$CUL1$^{R741E}$/RBX1; and 3: $^{N8}$CUL1$^{R717E}$/RBX1.

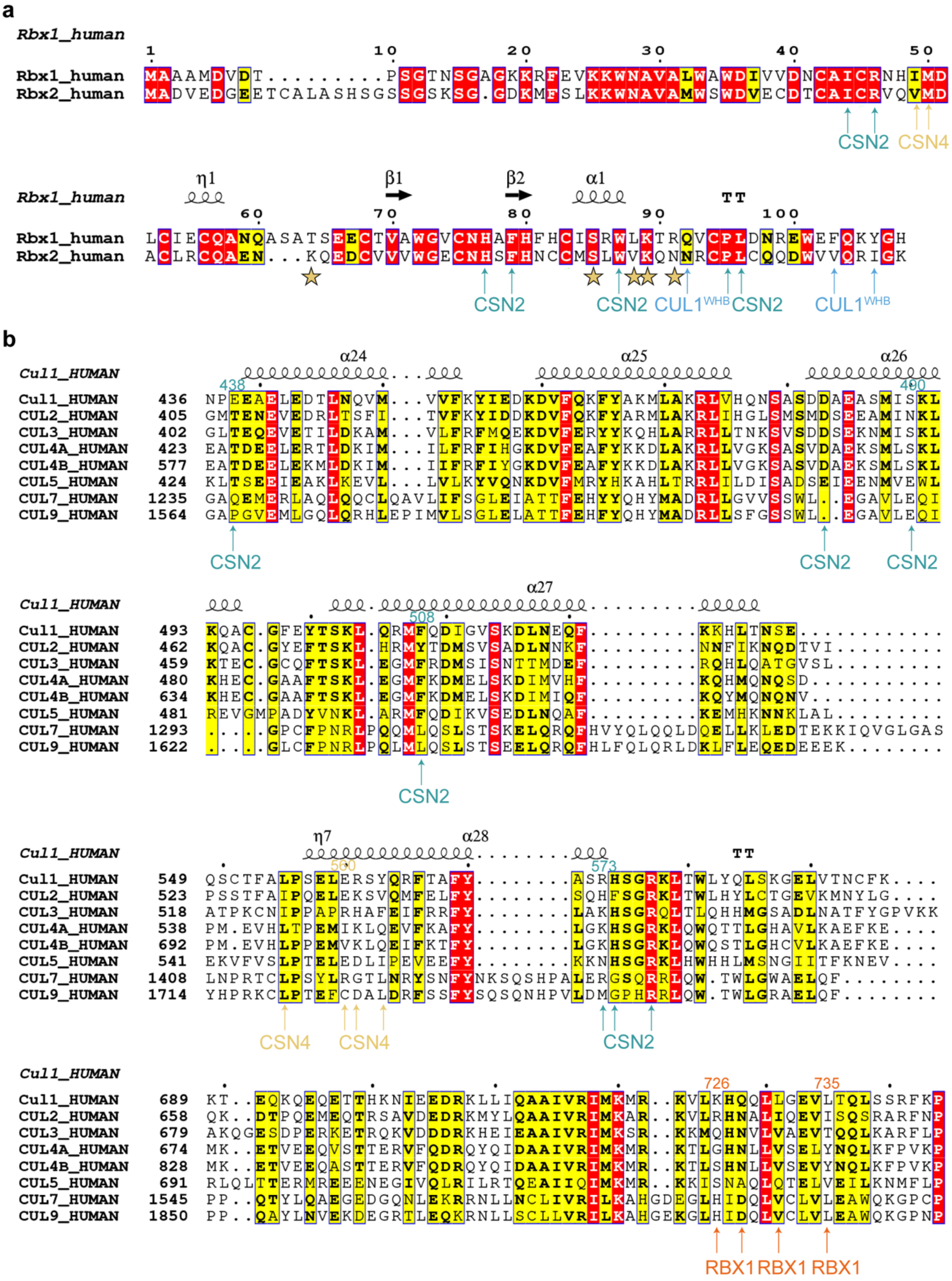
a
Rbx1_human
1 10 20 30 40 50
Rbx1_human MAAAMDVDT.........PSGTNSGAGKKRFEVKKWNAVALWAWDIVVDNCAICRNHIMD
Rbx2_human MADVEDGEETCALASHSGSSGSKSG.GDKMFSLKKWNAVAMWSWDVECDTCAICRVQVMD
CSN2 CSN4
η1 β1 β2 α1
Rbx1_human TT
60 70 80 90 100
Rbx1_human LCIECQANQASATSEECTVAWGVCNHAFHFHCISRWLKTRQVCPLDNREWEFQKYGH
Rbx2_human ACLRCQAEN...KQEDCVVVWGECNHSFHNCCMSLWVKQNNRCPLCQQDWVVQRIGK
CSN2 CSN2 CUL1WHB CSN2 CUL1WHB
b
α24 α25 α26
Cul1_HUMAN
438 490
Cul1_HUMAN 436 NPEEAELEDTLNQVM...VVFKYIEDKDVFQKFYAKMLAKRLVHQNSASDDAEASMISKL
CUL2_HUMAN 405 GMTENEVEDRLTSFI...TVFKYIDDKDVFQKFYARMLAKRLIHGLSMSMDSEEAMINKL
CUL3_HUMAN 402 GLTEQEVETILDKAM...VLFRFMQEKDVFERYYKQHLARRLLTNKSVSDDSEKNMISKL
CUL4A_HUMAN 423 EATDEELERTLDKIM...ILFRFIHGKDVFEAFYKKDLAKRLLVGKSASVDAEKSMLSKL
CUL4B_HUMAN 577 EATDEELEKMLDKIM...IIFRFIYGKDVFEAFYKKDLAKRLLVGKSASVDAEKSMLSKL
CUL5_HUMAN 424 KLTSEEIEAKLKEVL...LVLKYVQNKDVFMRYHKAHLTRRLILDISADSEIEENMVEWL
CUL7_HUMAN 1235 GAQEMERLAQLQQCLQAVLIFSGLEIATTFEHYYQHYMADRLLGVVSSWL..EGAVLEQI
CUL9_HUMAN 1564 GAPGVEMLGQLQRHLEPIMVLSGLELATTFEHFYQHYMADRLLSFGSSWL..EGAVLEQI
CSN2 CSN2 CSN2
α27
Cul1_HUMAN
508
Cul1_HUMAN 493 KQAC.GFEYTSKL.QRMFQDIGVSKDLNEQF.........KKHLTNSE............
CUL2_HUMAN 462 KQAC.GYEFTSKL.HRMYTDMSVSADLNNKF.........NNFIKNQDTVI.........
CUL3_HUMAN 459 KTEC.GCQFTSKL.EGMFRDMSISNTTMDEF.........RQHLQATGVSL.........
CUL4A_HUMAN 480 KHEC.GAAFTSKL.EGMFKDMELSKDIMVHF.........KQHMQNQSD...........
CUL4B_HUMAN 634 KHEC.GAAFTSKL.EGMFKDMELSKDIMIQF.........KQYMQNQNV...........
CUL5_HUMAN 481 REVGMPADYVNKL.ARMFQDIKVSEDLNQAF.........KEMHKNNKLAL.........
CUL7_HUMAN 1293 ....GPCFPNRLPQQMLQSLSTSKELQRQFHVYQLQQLDQELLKLEDTEKKIQVGLGAS
CUL9_HUMAN 1622 ....GLCFPNRLPQLMLQSLSTSEELQRQFHLFQLQRLDKLFLEQEDEEEK........
CSN2
η7 α28
Cul1_HUMAN TT
560 573
Cul1_HUMAN 549 QSCTFALPSELERSYQRFTAFY........ASRHSGRKLTWLYQLSKGELVTNCFK....
CUL2_HUMAN 523 PSSTFAIPQELEKSVQMFELFY........SQHFSGRKLTWLHYLCTGEVKMNYLG....
CUL3_HUMAN 518 ATPKCNIPPAPRHAFEIFRRFY........LAKHSGRQLTLQHHMGSADLNATFYGPVKK
CUL4A_HUMAN 538 PM.EVHLTPEMIKLQEVFKAFY........LGKHSGRKLQWQTTLGHAVLKAEFKE....
CUL4B_HUMAN 692 PM.EVHLPPEMVKLQEIFKTFY........LGKHSGRKLQWQSTLGHCVLKAEFKE....
CUL5_HUMAN 541 EKVFVSLPTELEDLIPEVEEFY........KKNHSGRKLHWHHLMSNGIITFKNEV....
CUL7_HUMAN 1408 LNPRTCLPSYLRGTLNRYSNFYNKSQSHPALERGSQRRLQW.TWLGWAELQF........
CUL9_HUMAN 1714 YHPRKCLPTEFCDALDRFSSFYSQSQNHPVLDMGPHRRLQW.TWLGRAELQF........
CSN4 CSN4 CSN2
Cul1_HUMAN
726 735
Cul1_HUMAN 689 KT..EQKQEQETTHKNIEEDRKLLIQAAIVRIMKMR..KVLKHQQLLGEVLTQLSSRFKP
CUL2_HUMAN 658 QK..DTPQEMEQTRSAVDEDRKMYLQAAIVRIMKAR..KVLRHNALIQEVISQSRARFNP
CUL3_HUMAN 679 AKQGESDPERKETRQKVDDDRKHEIEAAIVRIMKSR..KKMQHNVLVAEVTQQLKARFLP
CUL4A_HUMAN 674 MK..ETVEEQVSTTERVFQDRQYQIDAAIVRIMKMR..KTLGHNLLVSELYNQLKFPVKP
CUL4B_HUMAN 828 MK..ETVEEQASTTERVFQDRQYQIDAAIVRIMKMR..KTLSHNLLVSEVYNQLKFPVKP
CUL5_HUMAN 691 RLQLTTERMREEENEGIVQLRILRTQEAIIQIMKMR..KKISNAQLQTELVEILKNMFLP
CUL7_HUMAN 1545 PP..QTYLQAEGEDGQNLEKRRNLLNCLIVRILKAHGDEGLHIDQLVCLVLEAWQKGPCP
CUL9_HUMAN 1850 PP..QAYLNVEKDEGRTLEQKRNLLSCLLVRILKAHGEKGLHIDQLVCLVLEAWQKGPNP
RBX1 RBX1 RBX1

**Extended Data Fig. 10. Conservation of key interaction residues in CUL1 and RBX1.**

**a,** Sequence alignment of human RBX1 and RBX2, highlighting key residues involved in interactions with CSN2 (green arrows), CSN4 (yellow arrows) and $\mathrm{CUL}^{\mathrm{WHB}}$ (blue arrows). Residues specifically contributing to CSN4 interactions in dissociation-state-3 are marked with a yellow star. **b,** Sequence alignment of the human CUL1 family, with key residues mediating interactions with CSN2 (green arrows), CSN4 (yellow arrows) and RBX1 (orange arrows).

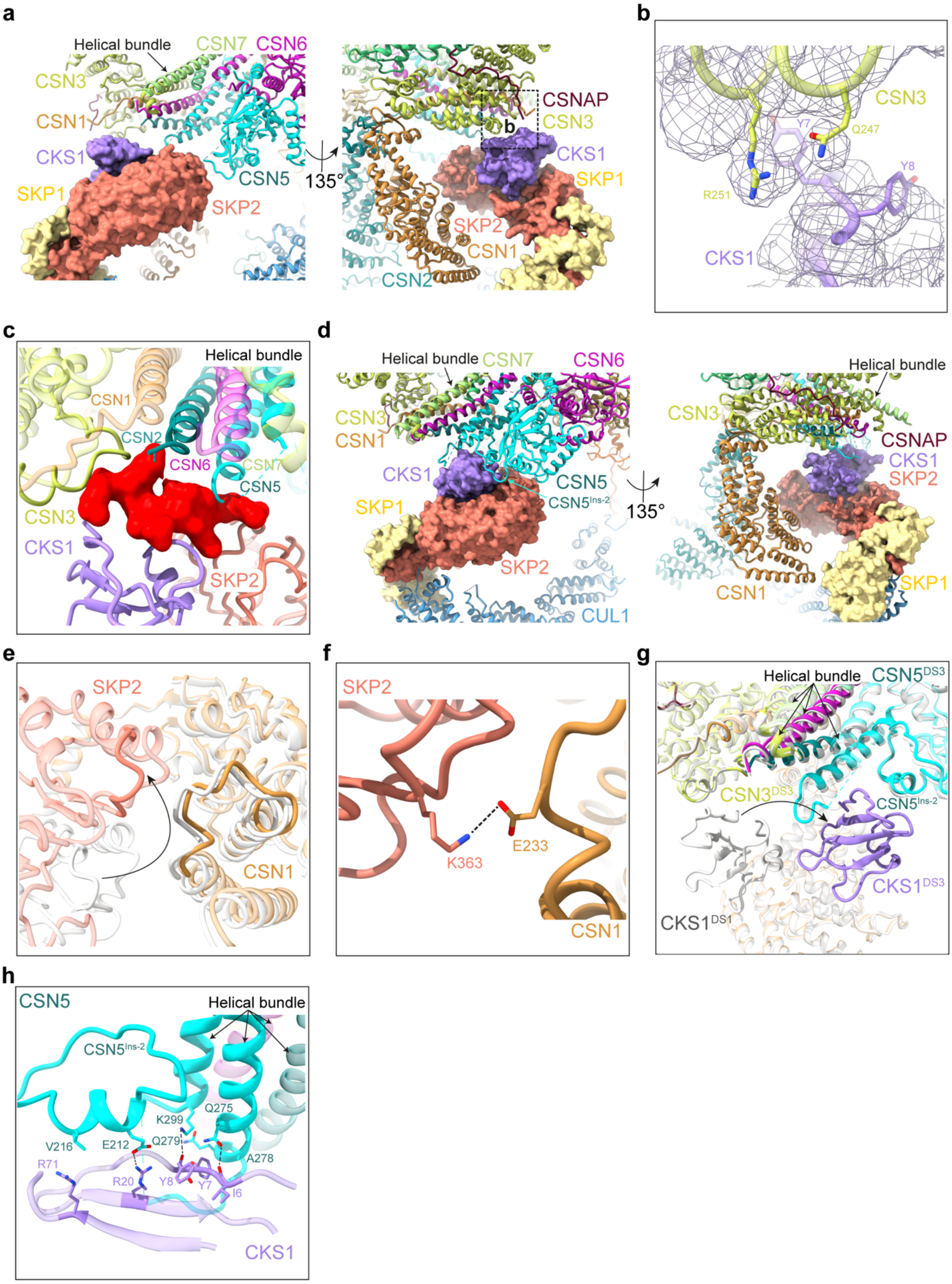

a
Helical bundle
CSN7
CSN6
CSN3
CSN1
CKS1
SKP1
CSN5
SKP2
135°
CSNAP
CSN3
b
CKS1
SKP1
SKP2
CSN1
CSN2
b
CSN3
Y7
Q247
Y8
R251
CKS1
c
Helical bundle
CSN1
CSN2
CSN6
CSN7
CSN5
CSN3
CKS1
SKP2
d
Helical bundle
CSN7
CSN6
CSN3
CSN1
CKS1
CSN5
CSN5Ins-2
SKP1
SKP2
CUL1
135°
Helical bundle
CSN3
CSNAP
CKS1
SKP2
SKP1
CSN1
e
SKP2
CSN1
f
SKP2
K363
E233
CSN1
g
Helical bundle
CSN5DS3
CSN3DS3
CSN5Ins-2
CKS1DS3
CKS1DS1
h
CSN5
Helical bundle
CSN5Ins-2
K299
Q275
V216
E212
Q279
A278
R71
R20
Y8
Y7
I6
CKS1

**Extended Data Fig. 11. Structural analysis of SR recognition by CSN.**

**a,** Location of the SR in pre-activated CSN$^{5H138A}$ -$^{N8}$SCF. SKP2 and CKS1 are positioned within a space formed by CSN1, CSN3, the CSN helical bundle, and CSN5. CSN subunits are displayed in ribbon, while the SR subunits are represented as a surface. **b,** In pre-activated CSN$^{5H138A}$ -$^{N8}$SCF, the SR subunit CKS1 appears to be stabilised by a small interface with CSN3, as supported by the cryo-EM density shown as a mesh. **c,** Docking of p27 (PDB: 2AST), the substrate of SKP1-CKS1, onto pre-activated CSN$^{5H138A}$ -$^{N8}$SCF, with p27 shown as a red surface. The model suggests that CSN cannot engage with the active SCF, as the presence of the substrate induces steric clashes with CSN. **d,** Location of the SR in CSN$^{E104A}$-SCF dissociation-state-3. Here, the SR occupies a space defined by the CSN1$^{arm}$, the CSN5 helical bundle, and the CSN5 MPN domain. CSN subunits are depicted in ribbon, while SR subunits are represented as a surface. **e,** Structural overlay highlighting key conformational changes in SKP2 between CSN$^{E104A}$-SCF dissociation-state-1 (grey) and CSN$^{E104A}$-SCF dissociation-state-3 (coloured), illustrating its structural rearrangement. **f,** Small but notable interface between SKP2 and CSN1 in CSN$^{E104A}$-SCF dissociation-state-3. **g,** Structural overlay highlighting key conformational changes in CKS1 between CSN$^{E104A}$-SCF dissociation-state-1 (grey) and CSN$^{E104A}$-SCF dissociation-state-3 (coloured). **h,** In CSN$^{E104A}$-SCF dissociation-state-3, there is an extensive interaction interface between CKS1, CSN5$^{Ins-2}$ and the CSN5 helical bundle.

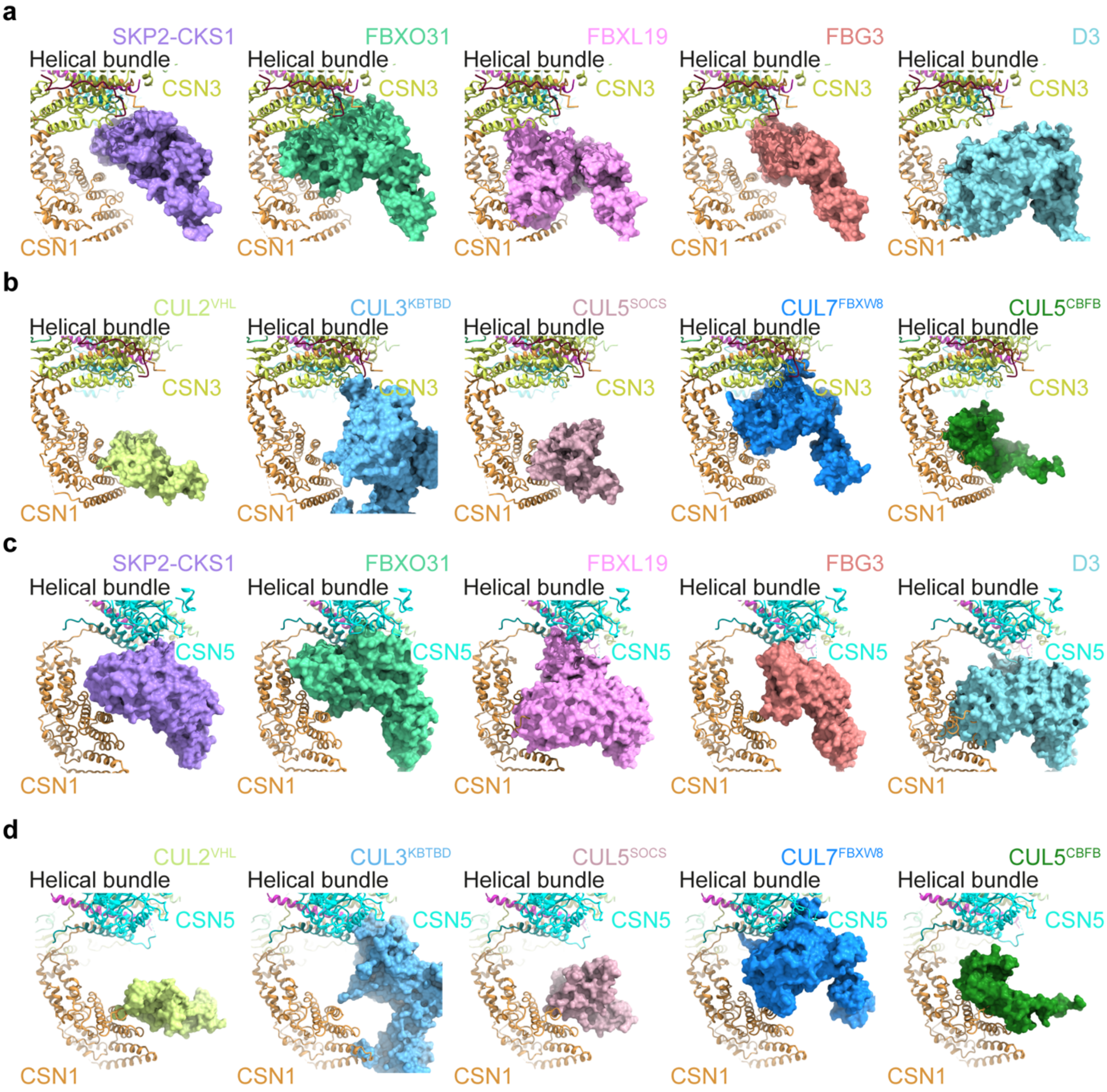


**Extended Data Fig. 12. Structural docking of diverse SRs onto pre-activated CSN$^{5H138A}$ -$^{N8}$SCF and CSN$^{E104A}$-SCF dissociation-state-3.**

**a,** Docking of various F-box SRs onto pre-activated CSN$^{5H138A}$ -$^{N8}$SCF reveals a conserved positioning, with the docked SRs aligning similarly to SKP2 in pre-activated CSN$^{5H138A}$ -$^{N8}$SCF. The PDB codes for the docked SRs are: FBXO31 (5VZT), FBXL19 (6WCQ), FBG3 (3WSO), and D3 (6BRO). **b**, Docking of SRs from different CRLs onto pre-activated CSN$^{5H138A}$ -$^{N8}$SCF demonstrates CSNs adaptability through an induced-fit mechanism, accommodating a broad range of SRs with distinct sizes and geometries. The PDB codes for the docked SRs are: CUL2$^{VHL}$ (4WQO), CUL3$^{KBTBD}$ (8H38), CUL5$^{SOCS}$ (4JGH), CUL7$^{FBXW8}$ (7Z8B), and CUL5$^{CBFB}$

(4N9F). **c,** Similar docking analysis of F-box protein SRs onto CSN$^{E104A}$-SCF dissociation-state-3 as described in (**a**). **d**, Docking of SRs from different CRLs onto CSN$^{E104A}$-SCF dissociation-state-3 as described in (**a**), highlights an additional role for CSN5 in SR recognition during later dissociation states.

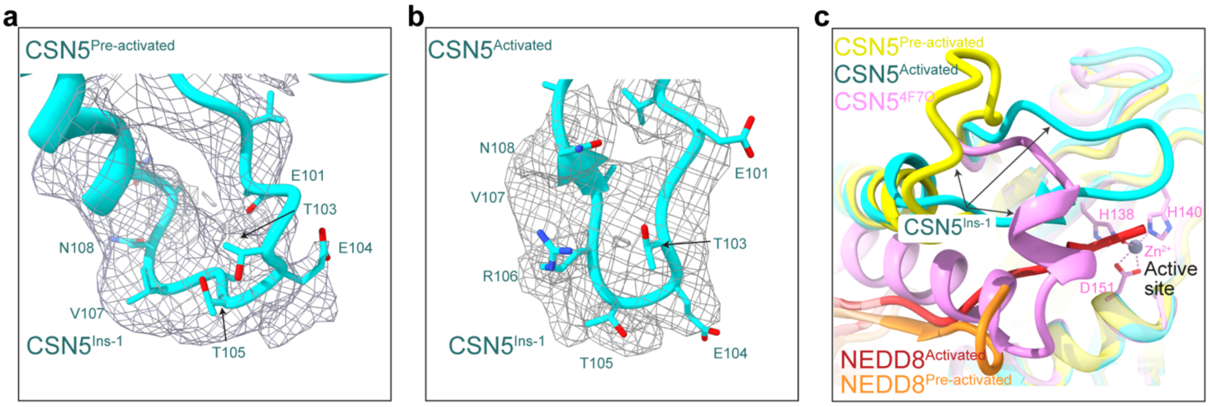


**Extended Data Fig. 13. Structural versatility of CSN5$^{Ins\text{-}1}$.**

**a,** Cryo-EM density and molecular model of CSN5$^{Ins\text{-}1}$ in pre-activated CSN$^{5H138A}$-$^{N8}$SCF. The side chains of CSN5$^{E101}$ and CSN5$^{E104}$ are modelled as approximations due to density limitations. **b,** Cryo-EM density and molecular model of CSN5$^{Ins\text{-}1}$ in activated CSN$^{5H138A}$-$^{N8}$SCF. **c,** Structural comparison of CSN5$^{Ins\text{-}1}$ from pre-activated and activated CSN$^{5H138A}$-$^{N8}$SCF, alongside the isolated crystal structure of CSN5 (PDB: 4F7O), demonstrating the conformational adaptability of this loop.

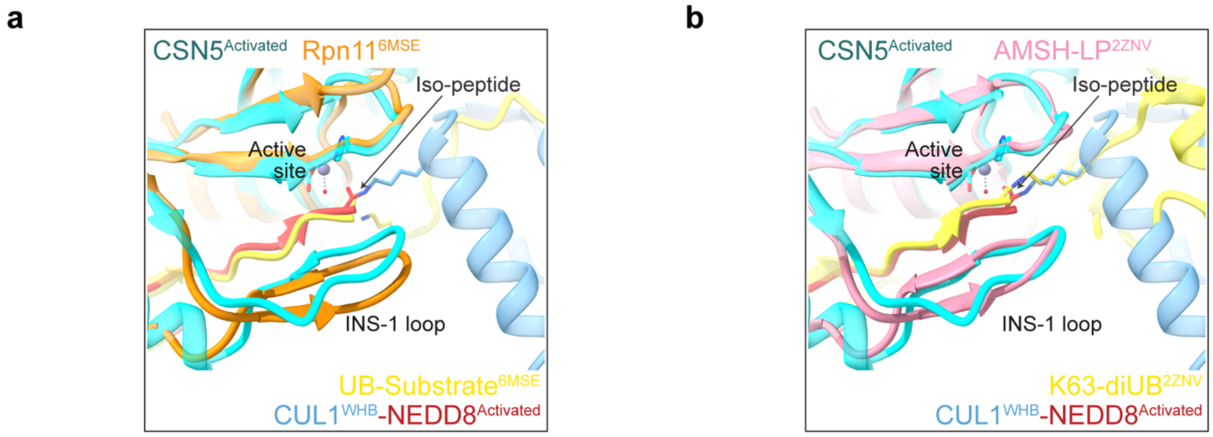


**Extended Data Fig. 14. Structural comparison of active sites in activated CSN$^{5H138A}$-$^{N8}$SCF and MPN$^{+}$ metalloprotease family members.**

**a,** Structural alignment of CSN5, NEDD8 and CUL1$^{WHB}$ in activated CSN$^{5H138A}$-$^{N8}$SCF with the RPN11-Ub from the 26S proteasome complex (PDB: 6MSE). **b,** Structural alignment of CSN5, and NEDD8 and CUL1$^{WHB}$ in activated CSN$^{5H138A}$-$^{N8}$SCF with the AMSH-LP-diUb complex (PDB: 2ZNV). In both cases, the conserved Ins-1 loop adopts a β-hairpin motif, stabilising the β-stranded C-terminal ubiquitin tail. However, in activated CSN$^{5H138A}$-$^{N8}$SCF, while CSN5$^{Ins\text{-}1}$ similarly stabilises the CUL1$^{K720}$-NEDD8$^{G76}$ isopeptide bond, it does not adopt a β-hairpin structure.

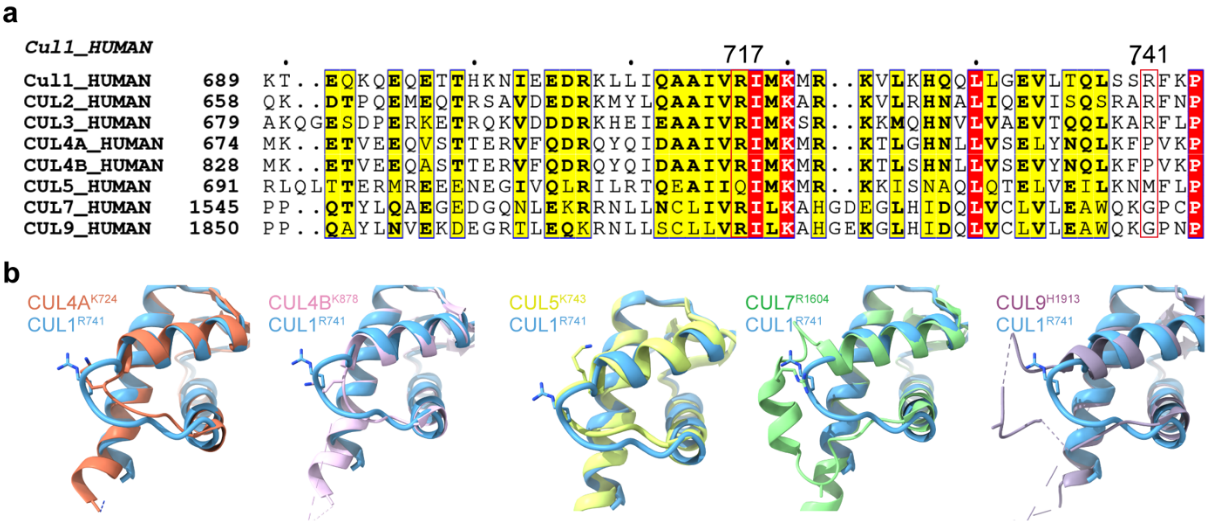


**Extended Data Fig. 15. Conservation Analysis of R717 and R741 across the cullin family.**

**a,** Sequence alignment of the human cullin family, highlighting $CUL1^{R717}$ and $CUL1^{R741}$ (red boxes). Sequence alignment was performed using the MUSCLE algorithm in Snapgene (Edgar 2004). **b,** Structural alignment of $CUL1^{R741}$ with CUL4A, CUL4B, CUL5, CUL7 and CUL9, demonstrating the structural conservation of $CUL1^{R741}$. PDB codes for respective cullins are as follows: CUL4A: 2HYE (Angers, Li et al. 2006); CULB: 4A0L (Fischer, Scrima et al. 2011); CUL5: 3DQV (Duda, Borg et al. 2008); CUL7: 7Z8B (Hopf, Baek et al. 2022); CUL9: 8Q7H (Horn-Ghetko, Hopf et al. 2024).

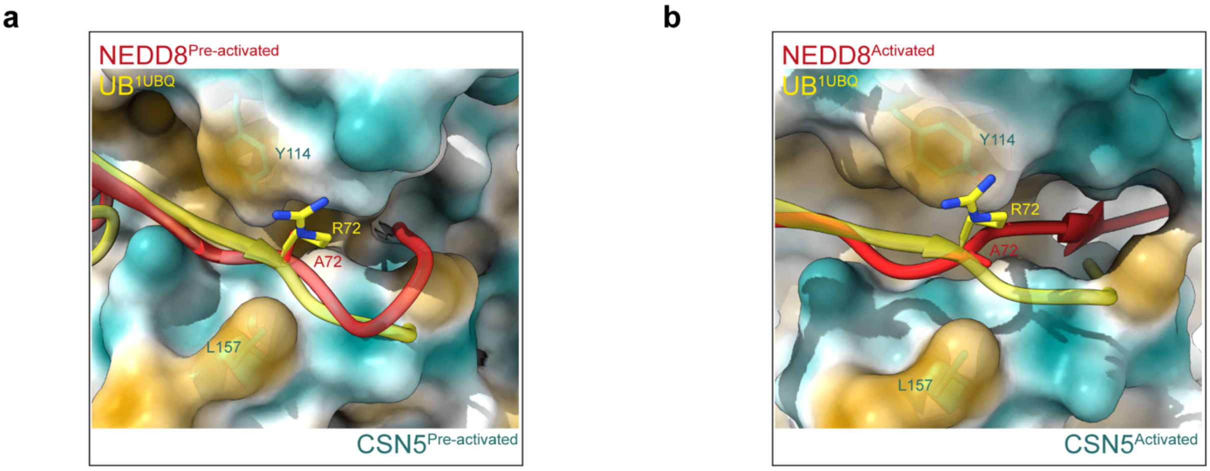


**Extended Data Fig. 16. CSN5 exhibits strict selectivity for NEDD8.**
**a,** Structural overlay of Ubiquitin (Ub) (PDB: 1UBQ) onto NEDD8 in pre-activated CSN$^{5H138A}$-$^{N8}$SCF. CSN5 is displayed as a hydrophobic surface. **b,** Structural overlay of Ubiquitin (PDB: 1UBQ) onto NEDD8 in activated CSN$^{5H138A}$-$^{N8}$SCF. The bulky charged Ub$^{R72}$ side chain is poorly accommodated within CSN5's hydrophobic pocket.

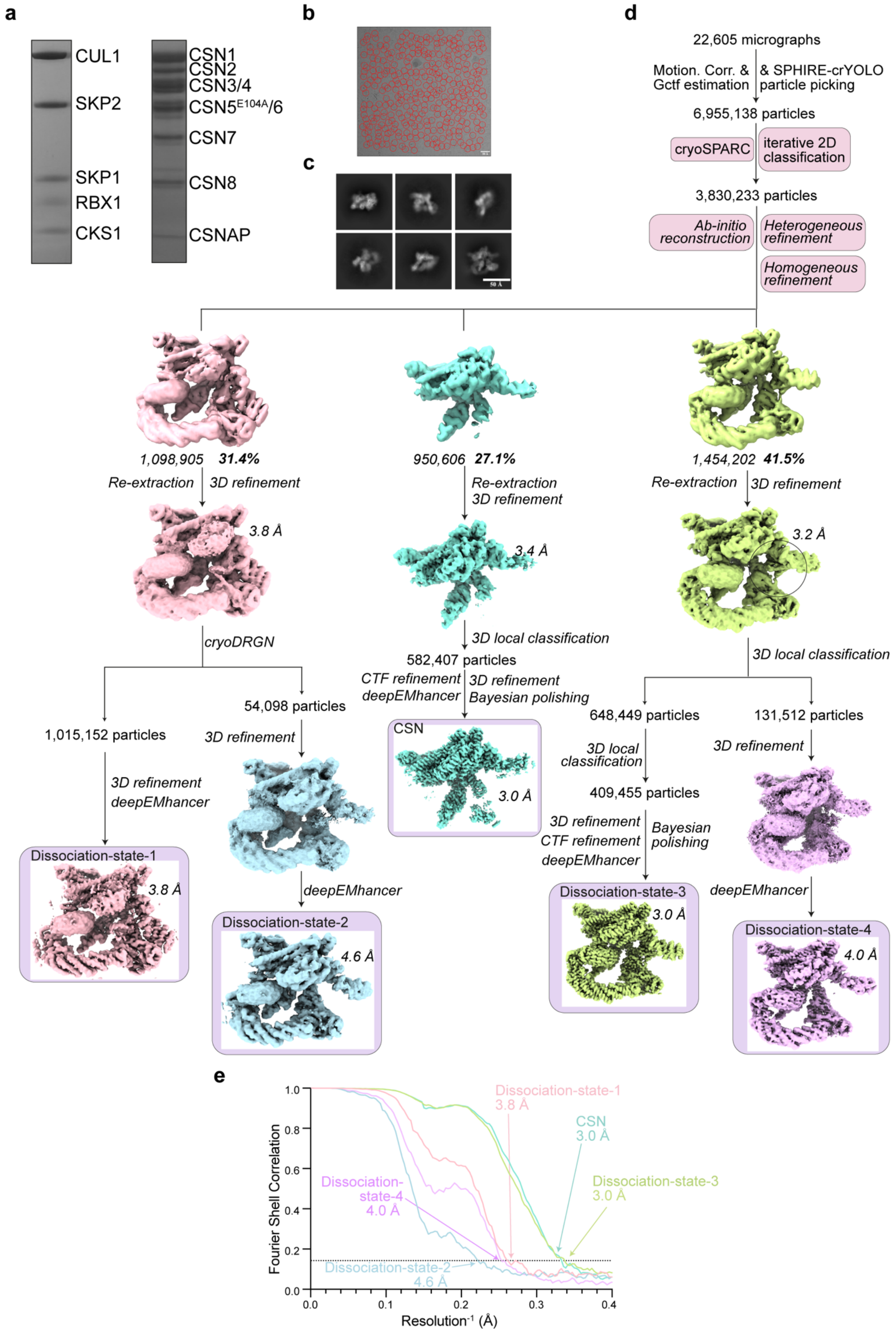
a
CUL1
SKP2
SKP1
RBX1
CKS1
CSN1
CSN2
CSN3/4
CSN5E104A/6
CSN7
CSN8
CSNAP
b
c
50 Å
d
22,605 micrographs
Motion. Corr. & Gctf estimation
& SPHIRE-crYOLO particle picking
6,955,138 particles
cryoSPARC
iterative 2D classification
3,830,233 particles
Ab-initio reconstruction
Heterogeneous refinement
Homogeneous refinement
1,098,905 31.4%
950,606 27.1%
1,454,202 41.5%
Re-extraction
3D refinement
3.8 Å
3.4 Å
3.2 Å
cryoDRGN
3D local classification
582,407 particles
CTF refinement
deepEMhancer
Bayesian polishing
CSN
3.0 Å
54,098 particles
1,015,152 particles
648,449 particles
131,512 particles
409,455 particles
Dissociation-state-1
Dissociation-state-2
4.6 Å
Dissociation-state-3
Dissociation-state-4
4.0 Å
e
Fourier Shell Correlation
Resolution-1 (Å)
Dissociation-state-1 3.8 Å
CSN 3.0 Å
Dissociation-state-3 3.0 Å
Dissociation-state-4 4.0 Å
Dissociation-state-2 4.6 Å

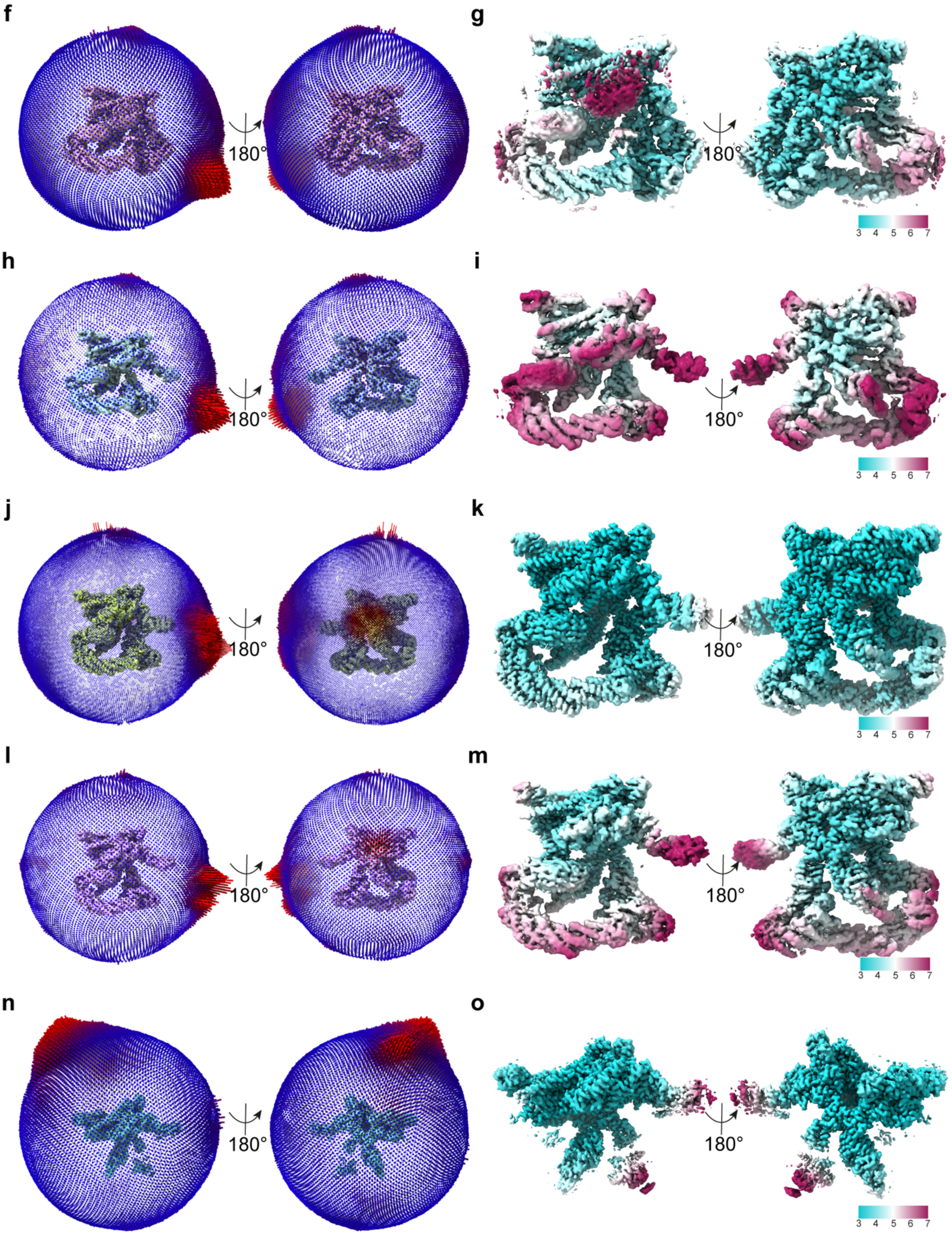


**Extended Data Fig. 17. Cryo-EM and single particle analysis on CSN$^{5E104A}$-SCF dissociation states.**

**a,** SDS-PAGE (4-12%) analysis and Coomassie staining of assembled SCF complex (CUL1, RBX1, SKP1, SKP2 and CKS1) and reconstituted CSN5$^{E104A}$. **b,** A representative motion-corrected micrograph and particle picking (red circles). **c,** Representative high-quality 2D reference-free 2D class averages. **d,** Single particle analysis workflow for the CSN$^{5E104A}$-SCF dataset. 3D classifications were performed in CryoDRGN (Zhong, Bepler et al. 2021) and RELION-4.0 (Kimanius, Dong et al. 2021)). **e,** Resolution estimates of maps resolved from the CSN$^{E104A}$-SCF dataset. **f,** Euler angle distribution plots for dissociation-state-1. **g,** Local resolution estimates for dissociation-state-1. **h,** Euler angle distribution plots for dissociation-state-2. **i,** Local resolution estimates of dissociation-state-2. **j,** Euler angle distribution plots for dissociation-state-3. **k,** Local resolution estimates of dissociation-state-3. **l,** Euler angle distribution plots of dissociation-state-4. **m,** Local resolution estimates of dissociation-state-1. **n,** Euler angle distribution plots of free CSN (CSN$^{Apo}$). **o,** Local resolution estimates of free CSN (CSN$^{Apo}$). Resolutions for all maps in this figure were estimated using the gold-standard FSC 0.143 criterion.

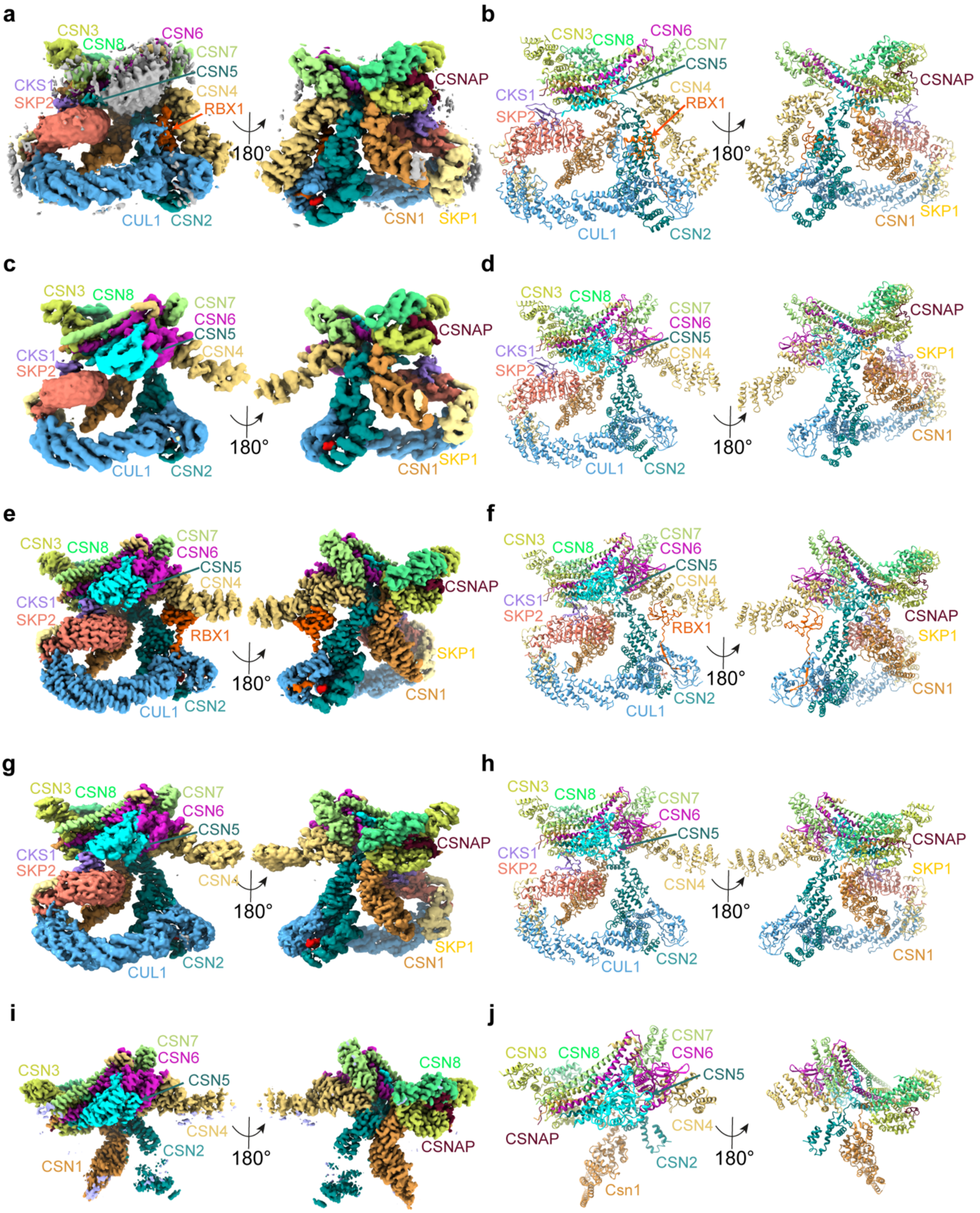


**Extended Data Fig. 18. Cryo-EM structures from CSN$^{E104A}$-SCF dissociation states.**

**a,** Cryo-EM density map of dissociation-state-1. **b,** Molecular model of dissociation-state-1 (ribbon representation). **c,** Cryo-EM density map of dissociation-state-2. **d,** Molecular model of dissociation-state-2 (ribbon representation). **e,** Cryo-EM density map of dissociation-state-3. **f,** Molecular model of dissociation-state-3 (ribbon representation). **g,** Cryo-EM density map of dissociation-state-4. **h,** Molecular model of dissociation-state-4 (ribbon representation). **i,** Cryo-EM density map of fully dissociated $CSN^{apo}$. **j,** Molecular model of fully dissociated $CSN^{apo}$ (ribbon representation).

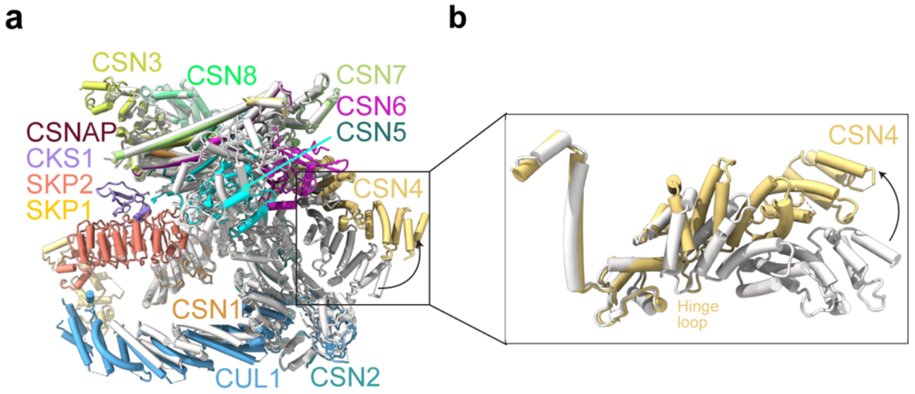


**Extended Data Fig. 19. Structural comparison between activated $CSN^{5H138A}$-$^{N8}$SCF and $CSN^{E104A}$-SCF dissociation-state-2.**

**a,** Structural overlay highlighting conformational changes in CSN between activated $CSN^{5H138A}$-$^{N8}$SCF (grey) and $CSN^{E104A}$-SCF dissociation-state-2 (coloured). **b,** Close-up view of the boxed region in (**a**), illustrating the repositioning of the $CSN4^{arm}$ about the $CSN4^{hinge\text{-}loop}$.

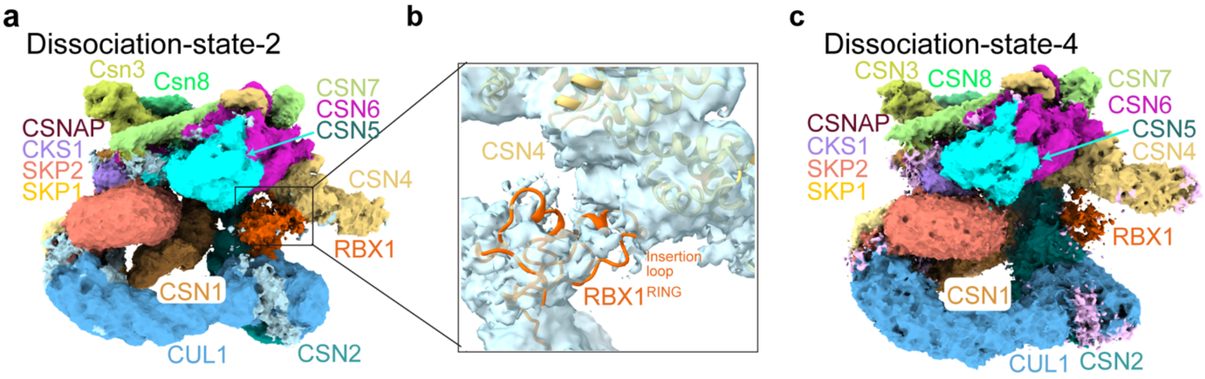


**Extended Data Fig. 20. Visualisation of RBX1$^{RING}$ in unprocessed cryo-EM maps of CSN$^{E104A}$-SCF dissociation-state-2 and dissociation-state-4.**

**a,** Final RELION-4.0 refined cryo-EM map (Kimanius, Dong et al. 2021) of CSN$^{E104A}$-SCF dissociation-state-2, revealing weak but distinct density for RBX1$^{RING}$. **b,** Close-up view of the boxed region in (**a**), featuring a "best-fit" molecular model for RBX1 within the observed density. Notably, a potential contact is evident between the RBX1$^{RING\text{-}Insertion}$ and the CSN4$^{arm}$. **c,** Final RELION-4.0 refined map of CSN$^{E104A}$-SCF dissociation-state-4, where only faint density corresponding to a portion of RBX1$^{RING}$ is detectable at very low contour levels.

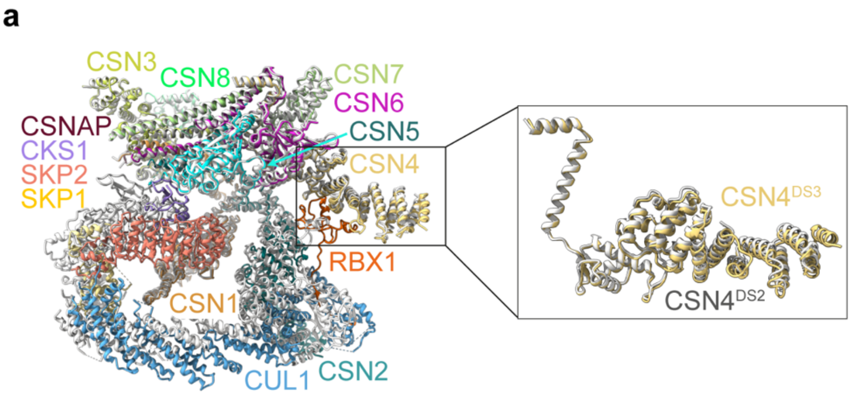


**Extended Data Fig. 21. Structural comparison between CSN$^{E104A}$-SCF dissociation-state-2 and dissociation-state-3.**

**a,** Structural overlay of CSN$^{E104A}$-SCF dissociation-state-2 (grey) and dissociation-state-3 (coloured). A close-up view of the boxed region reveals that the CSN4$^{arm}$ adopts a similar positioning in both states.

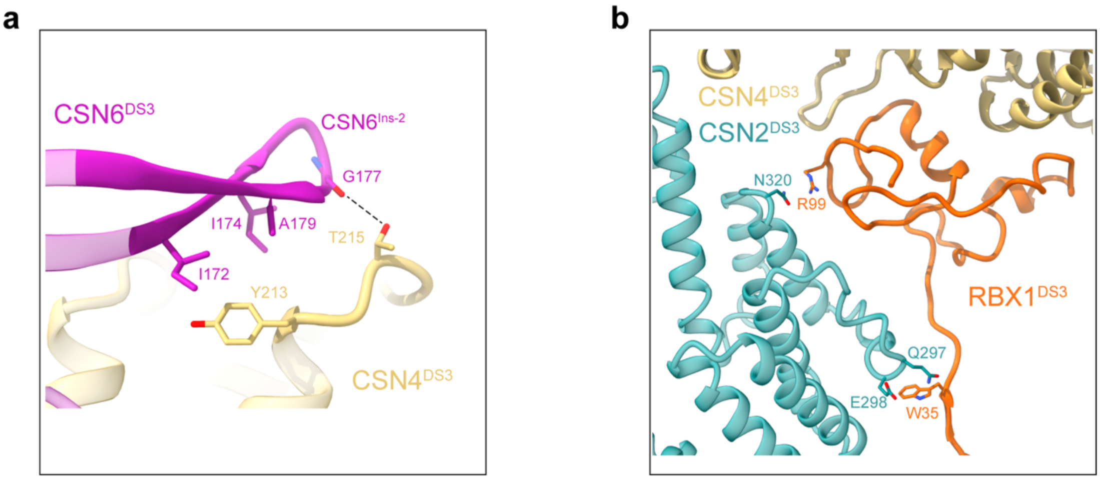


**Extended Data Fig. 22. Key interfaces describing CSN$^{E104A}$-SCF dissociation-state-3.**

**a,** CSN6$^{Ins\text{-}2}$ interacts with CSN4, securing the CSN5-CSN6 MPN domains in an autoinhibited conformation. **b,** The interface between CSN2 and RBX1 in dissociation-state-3 is minimal, indicating a reduced stabilising interaction compared to activated CSN$^{5H138A}$-$^{N8}$SCF.

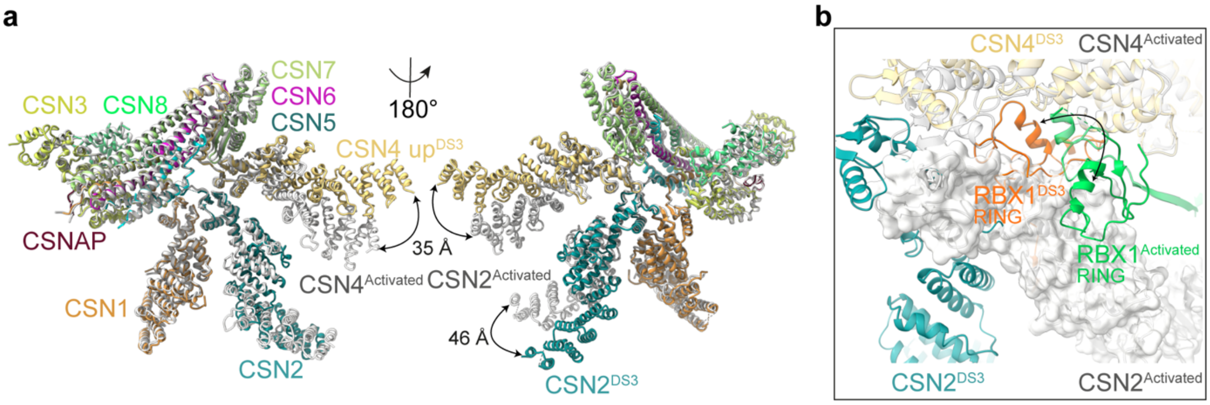


**Extended Data Fig. 23. Conformational changes between activated $CSN^{5H138A}$-$^{N8}SCF$ and $CSN^{E104A}$-SCF dissociation-state-3.**

**a,** Structural overlay highlighting key conformational changes in $CSN2^{arm}$ and $CSN4^{arm}$ between activated $CSN^{5H138A}$-$^{N8}SCF$ (grey) and $CSN^{E104A}$-SCF dissociation-state-3 (coloured). **b,** Remodelling of $RBX1^{RING}$ from its position in activated $CSN^{5H138A}$-$^{N8}SCF$ (green) to $CSN^{E104A}$-SCF dissociation-state-3 (orange). In dissociation-state-3, the repositioned $RBX1^{RING}$ results in a significant steric clash with $CSN2^{arm}$ in activated $CSN^{5H138A}$-$^{N8}SCF$ (grey surface). The structural alignment was performed using pre-activated CSN4 as a reference.

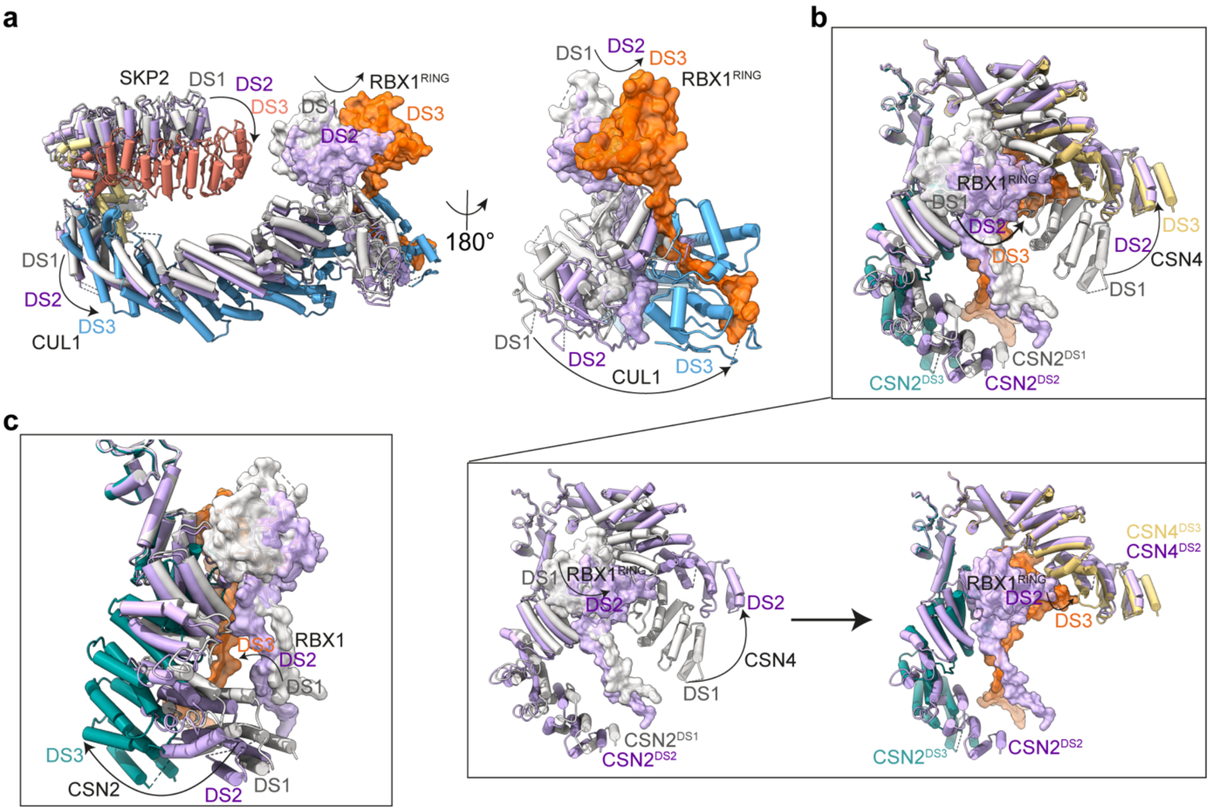


**Extended Data Fig. 24. Stepwise conformational changes in SCF, CSN2$^{arm}$ and CSN4$^{arm}$ during CSN$^{E104A}$-SCF dissociation.**

**a,** Progressive structural transitions within **SCF** across dissociation-state-1 (grey), dissociation-state-2 (purple) and dissociation-state-3 (colour scheme used in previous figures). SCF subunits are depicted in ribbon (tube format) with RBX1 highlighted by an additional semi-transparent surface. The RBX1$^{RING}$ in dissociation-state-2 is a best-fit model for illustrative purposes only. **b,** Stepwise conformational rearrangements of CSN4$^{arm}$ relative to RBX1$^{RING}$. Top panel: overlay of sequential transitions from dissociation-states-1, -2 and –3. Bottom panel: distinct transitions from dissociation-state-1 (grey) to dissociation-state-2 (purple), followed by dissociation-state-2 (purple) to dissociation-state-3 (colour scheme used in previous figures). **c,** Sequential conformational changes of CSN2$^{arm}$ relative to RBX1, illustrating structural movements across dissociation-states-1, -2 and -3.

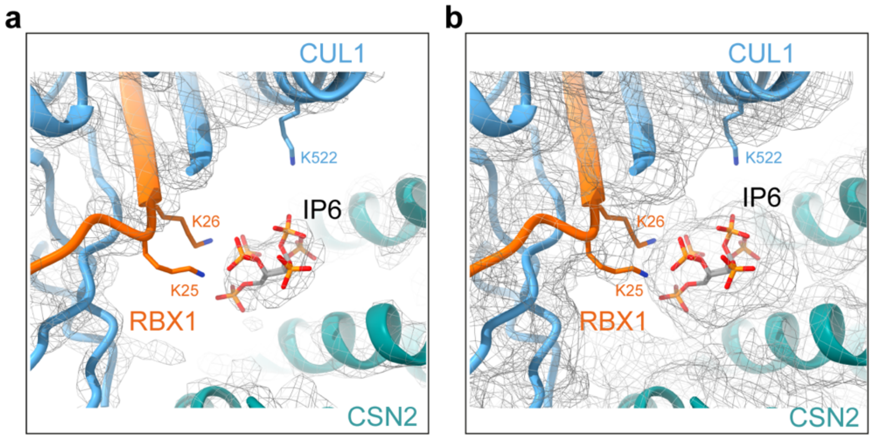


**Extended Data Fig. 25. CSN2-bound IP6 and its interactions with CUL1/RBX1 in $CSN^{E104A}$-SCF dissociation-state-3.**

**a,** Cryo-EM density map from $CSN^{E104A}$-SCF dissociation-state-3, highlighting the density for IP6 bound to CSN2. **b,** Low-contour view of the same map in (**a**), revealing side chain densities for $RBX1^{K25}$, $RBX1^{K26}$ and $CUL1^{K522}$, suggesting potential interaction sites between IP6 and CUL1/RBX1 in dissociation-state-3.

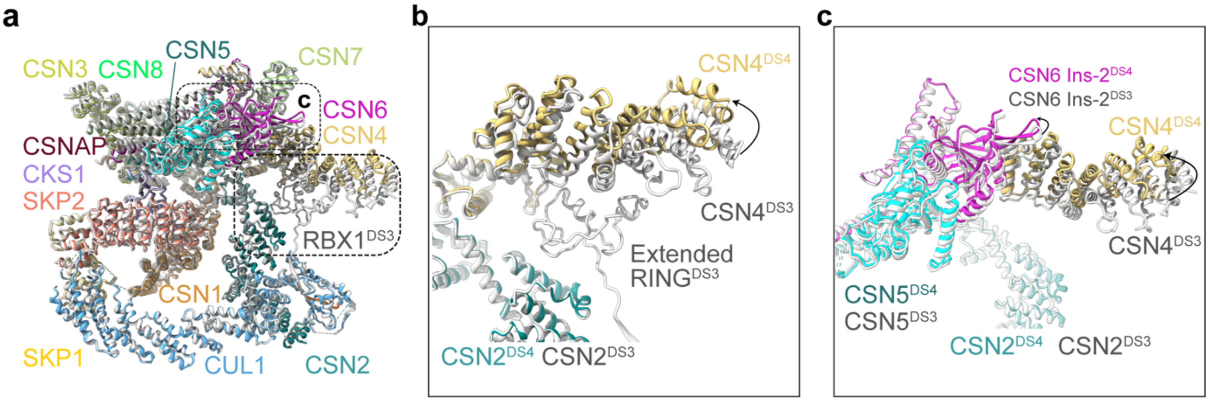


**Extended Data Fig. 26. Structural comparison between CSN$^{E104A}$-SCF dissociation-state-3 and dissociation-state-4.**

**a,** Structural overlay highlighting key conformational changes in CSN between CSN$^{E104A}$-SCF dissociation-state-3 (grey) and CSN$^{E104A}$-SCF dissociation-state-4 (coloured). **b,** Close-up view of the boxed region in (**a**), illustrating an upward shift of the CSN4$^{arm}$ in dissociation-state-4, which disrupts the RBX1$^{RING}$ interface observed in dissociation-state-3. **c,** Close-up view of the boxed region in (**a**), showing the repositioning of CSN6$^{Ins\text{-}2}$ in dissociation-state-4, which maintains its interface with CSN4$^{arm}$.

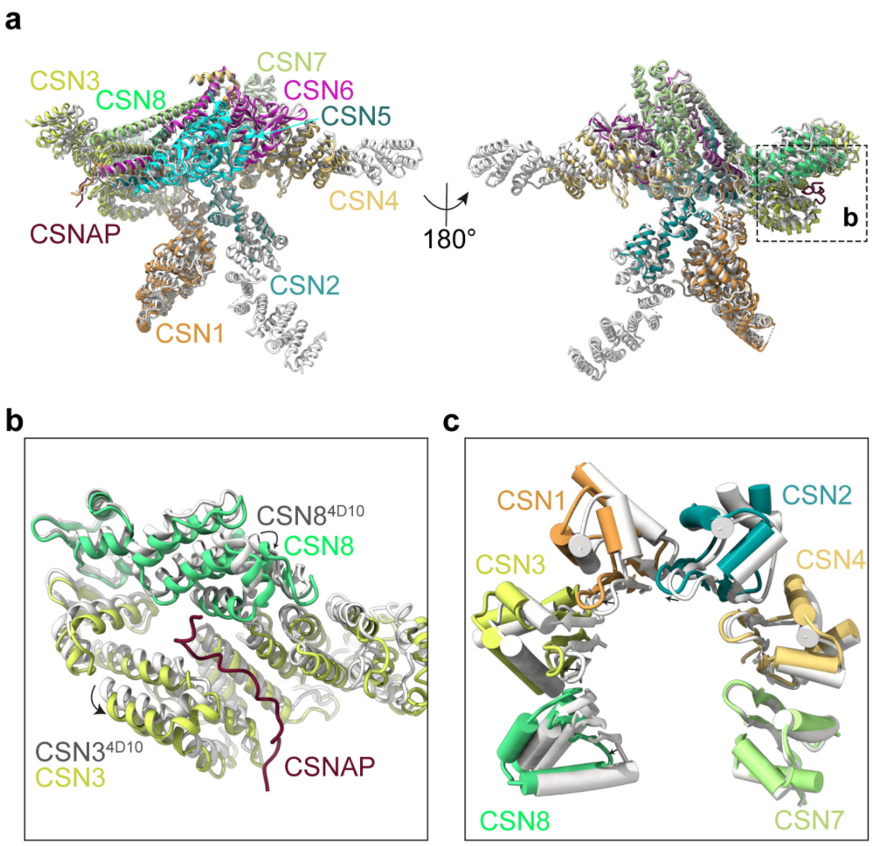


**Extended Data Fig. 27. Structural rearrangements in CSN upon CSNAP incorporation.**

**a,** Structural overlay of the 9-subunit CSN complex (coloured) and the 8-subunit CSN complex (PDB: 4D10) (grey). **b,** Close-up view of the boxed region in (**a**), illustrating structural adjustments in CSN3 and CSN8 upon CSNAP incorporation. **c,** Conformational changes within the PCI ring upon incorporation of CSNAP.

Angers, S., T. Li, X. Yi, M. J. MacCoss, R. T. Moon and N. Zheng (2006). "Molecular architecture and assembly of the DDB1-CUL4A ubiquitin ligase machinery." Nature **443**(7111): 590-593.

Duda, D. M., L. A. Borg, D. C. Scott, H. W. Hunt, M. Hammel and B. A. Schulman (2008). "Structural insights into NEDD8 activation of cullin-RING ligases: conformational control of conjugation." Cell **134**(6): 995-1006.

Edgar, R. C. (2004). "MUSCLE: multiple sequence alignment with high accuracy and high throughput." Nucleic Acids Res **32**(5): 1792-1797.

Fischer, E. S., A. Scrima, K. Bohm, S. Matsumoto, G. M. Lingaraju, M. Faty, T. Yasuda, S. Cavadini, M. Wakasugi, F. Hanaoka, S. Iwai, H. Gut, K. Sugasawa and N. H. Thoma (2011). "The molecular basis of CRL4DDB2/CSA ubiquitin ligase architecture, targeting, and activation." Cell **147**(5): 1024-1039.

Hopf, L. V. M., K. Baek, M. Klugel, S. von Gronau, Y. Xiong and B. A. Schulman (2022). "Structure of CRL7(FBXW8) reveals coupling with CUL1-RBX1/ROC1 for multi-cullin-RING E3-catalyzed ubiquitin ligation." Nat Struct Mol Biol **29**(9): 854-862.

Horn-Ghetko, D., L. V. M. Hopf, I. Tripathi-Giesgen, J. Du, S. Kostrhon, D. T. Vu, V. Beier, B. Steigenberger, J. R. Prabu, L. Stier, E. M. Bruss, M. Mann, Y. Xiong and B. A. Schulman (2024). "Noncanonical assembly, neddylation and chimeric cullin-RING/RBR ubiquitylation by the 1.8 MDa CUL9 E3 ligase complex." Nat Struct Mol Biol **31**(7): 1083-1094.

Kimanius, D., L. Dong, G. Sharov, T. Nakane and S. H. W. Scheres (2021). "New tools for automated cryo-EM single-particle analysis in RELION-4.0." Biochem J **478**(24): 4169-4185.

Zhong, E. D., T. Bepler, B. Berger and J. H. Davis (2021). "CryoDRGN: reconstruction of heterogeneous cryo-EM structures using neural networks." Nat Methods **18**(2): 176-185.
